# Selective inhibition in CA3: A mechanism for stable pattern completion through heterosynaptic plasticity

**DOI:** 10.1101/2024.08.16.608240

**Authors:** Gyeongtae Kim, Pilwon Kim

## Abstract

Neural assemblies representing different engrams compete for successful retrieval in the CA3 region of the hippocampus, yet the detailed mechanisms underlying their formation remain elusive. Recent research indicates that hippocampal inhibitory neurons respond selectively to stimuli and exhibit diverse plasticity, suggesting their significant role in engram formation. Conventional attractor network models for CA3 typically employ global inhibition, where inhibitory neurons uniformly suppress the activity of excitatory neurons. However, these models may not fully capture the complex dynamics of competition arising from sparse distributed coding and may not accurately reflect the specific roles of inhibitory neurons in the competition between neural assemblies during memory retrieval. We propose a mechanism for engram formation in CA3 using a spiking neural network model, emphasizing the critical role of the association between excitatory and inhibitory neurons through heterosynaptic plasticity. In our model, inhibitory neurons are associated with specific neural assemblies during encoding and selectively inhibit excitatory neurons involved in competing assemblies during retrieval. With a simplified dentate gyrus (DG) in a feed-forward structure, this proposed mechanism results in sparsely distributed engrams in CA3. The sparse distributed coding in the model allows us to investigate the effects of selective inhibition on pattern completion under various configurations, such as partially overlapping competing engrams. Our results demonstrate that selective inhibition provides more stable pattern completion and enhances retrieval performance compared to global inhibition alone. Furthermore, the observed neural activity in the hippocampal subregions of the model aligns with experimental findings on these regions’ roles in pattern separation and pattern completion.

**Author Summary:** We explored how memories are stored and retrieved in the hippocampus by focusing on the CA3 region, a critical component of memory processes. Using a spiking neural network model, we propose a new mechanism in which specific inhibitory neurons selectively control the activity of other neurons during memory retrieval. We found that this selective inhibition can be naturally induced during memory encoding. This selective inhibition offers an alternative to traditional models that assume global suppression and provides a more nuanced understanding of how memories compete for successful retrieval. Our findings suggest that this selective inhibition improves the stability and accuracy of memory recall. The model also aligns with known biological functions of the hippocampus, shedding light on how complex memory processes, such as distinguishing between similar memories and accurately reconstructing past experiences, might be managed in the brain. This research offers new insights into the dynamic roles of inhibitory neurons in balancing memory encoding and retrieval, enhancing our understanding of memory function.

## 2 Introduction

The hippocampus is a crucial region for processing episodic memory (1; 2). In this region, engrams—neural circuits representing the physical traces of memories—are stored and retrieved (3; 4). Engrams are specifically formed in CA3 in a sparse and distributed manner, facilitated by sparse activations from the DG (5). This process, known as pattern separation, creates distinct and non-overlapping representations of similar events or stimuli. It is one of the key functions of the hippocampus, which reduces interference between memories (6; 7; 8; 9; 10). Successful pattern separation is achieved when distinct representations in CA3 are formed regardless of similarities in external information (11; 12; 13; 14).

While the DG generates sparse and highly distinct representations that substantially differ from the original inputs, the hippocampus can still successfully retrieve specific memories through CA3. This process, the reconstruction of a complete memory from partial or incomplete cues, is called pattern completion (6; 7; 15; 16; 9; 10; 17). It relies on two critical hippocampal pathways: the direct perforant path (PP) from the entorhinal cortex (EC) to CA3, and the recurrent collateral (Rc) connections within CA3. The direct PP encodes features that enable neural assemblies in CA3 to recognize similarities between incoming cues and stored memories. As an attractor network, CA3 uses Rc connections to activate the full neural assembly, stabilizing the network and ensuring accurate memory retrieval (18; 19; 20; 21). These Rc connections specifically enhance the activity of relevant neurons, fully reactivating the correct memory trace while suppressing interference from competing memories.

Although the processes of pattern separation and pattern completion in the hippocampus are well understood, the underlying mechanisms related to engrams in CA3 remain elusive, particularly concerning the role of inhibitory neurons in these processes. Attractor network models describe pattern completion in CA3 in the context of memory competition (18; 19; 20; 21), but they do not specifically address the mechanisms underlying engram formation. Most of these models assume completely separate neural assemblies that compete through a winner-takes-all process. They commonly employ global inhibition, where inhibitory neurons suppress all excitatory neurons.

Recent research has highlighted the crucial role of inhibitory neurons in the memory process (22; 23). Stimulus selectivity was primarily associated with excitatory neurons (24; 25). However, recent studies reveal that many GABAergic neurons exhibit greater stimulus selectivity than previously recognized.(26; 27). In addition, increasing evidence shows the presence of synaptic plasticity mechanisms in inhibitory neurons. Both long-term potentiation (LTP) and long-term depression (LTD) have been observed at inhibitory neuron synapses, similar to what has been previously observed at excitatory synapses (28; 29; 30; 31; 32). This plasticity contributes to engram formation and refinement by modulating inhibitory control in response to experience. Extending beyond their contribution to sparse distributed coding (33; 34), these observations suggest that inhibitory neurons play a broader role in memory encoding, storage, and retrieval processes.

Building on the evidence discussed above, our work aims to explore the roles of inhibitory neurons in memory processing using a modeling approach. We propose that inhibitory neurons offer functional advantages by providing selective inhibition in memory competition, beyond the effects of global inhibition, potentially enhancing the stability and efficiency of memory retrieval. We developed a spiking neural network model that preserves the structural and connectional properties of the hippocampus, with a focus on the trisynaptic circuit—the pathway from the EC to CA3 via the DG. Drawing on evidence of heterosynaptic plasticity observed in various connections (35; 36; 37; 38), we propose a mechanism of engram formation where heterosynaptic plasticity between excitatory and inhibitory neurons in CA3 naturally induces selective inhibition during memory competition.

In the proposed model, selective inhibition is characterized by inhibitory neurons that are associated with a specific neural assembly in CA3: these neurons inhibit excitatory neurons in competing assemblies while sparing those within their own assembly. To investigate the effect of selective inhibition on pattern completion, we simplified the DG into a feed-forward structure. This simplification enables instant sparse distributed coding and, together with the proposed engram formation mechanism, creates various competitive conditions. These conditions result in situations within CA3 where multiple overlapping engrams of varying sizes compete during memory retrieval.

We conducted pattern separation and pattern completion tasks across the entire hippocampal model, made possible by the stable retrieval performance. The resulting activation patterns in each subregion were then compared with experimental data. The analysis shows that selective inhibition by inhibitory neurons enhances the stability and efficiency of competition between neural assemblies. These findings provide novel insights into the role of inhibitory neurons in pattern completion and separation within the hippocampus.

## 3 Results

### 3.1 Model architecture and learning mechanisms

We constructed a spiking neural network that preserves hippocampal structure to model engram formation in CA3 and the dynamics of engram competition during retrieval (‘Materials and methods’). The primary goal of this model is to evaluate the stability of pattern completion across a range of conditions. To achieve this, engrams in CA3 should be naturally formed for arbitrary inputs via the DG, resulting in a sparse distributed representation. However, modeling DG pattern separation is challenging due to its intricate circuitry, complex synaptic connectivity, and heterogeneous neuronal types (39; 40; 41). Therefore, we adopted a simplified approach by focusing solely on spatial pattern separation for non-sequential inputs, represented as 16-dimensional binary vectors at a fixed frequency (‘Materials and methods’). Under these conditions, we designed a feedforward form of the DG, which we will explain in more detail in the next section, ‘Feedforward form of the dentate gyrus and its inherent ability for spatial pattern separation.’

Additionally, we simplified CA1’s role to reinstating original information from CA3 engrams (‘Materials and Methods’). All neurons in the network were modeled using the Izhichevich model (42) distinguishing hippocampal neurons as either excitatory or inhibitory (‘Materials and Methods’). The model operates in distinct phases according to theta oscillation theory (43). Theta oscillations (4-8 Hz) are prominent in the hippocampus and are crucial in memory processes. Several theories have been proposed to explain the specific functions of theta waves (43; 44; 45). One theory suggests that theta oscillations divide information processing into discrete time windows, optimizing memory by coordinating information flow across hippocampal subregions (46; 47; 48; 49; 50; 51; 52; 53).

Based on theta oscillation theory, during the encoding phase, superficial EC sends inputs to both DG and deep EC, with DG relaying signals to CA3 and deep EC forwarding inputs to CA1 (Fig 1A1). In CA3, four primary connections crucial for retrieval-direct PP, Rc, Schaffer collateral (Sc), and the connection from CA1 to deep EC—are silenced during encoding (except for inhibitory connections within CA3; see Table 3). during the encoding. This phase-specific silencing prevents interference from previously learned memories, thereby maintaining the integrity of stored information while allowing new patterns to be encoded efficiently. Despite being silenced, the direct PP, Sc, and excitatory Rc connections within CA3 continue to undergo learning via heterosynaptic plasticity, as described in the next section (Fig 1C, and Table 3). To simplify the model, we applied symmetric spike-timing-dependent plasticity (STDP) across all connections, thereby unifying the different forms of STDP observed in various pathways, based on symmetric STDP identified in the CA3 recurrent circuit (‘Materials and Methods’ and Fig 1B) (54; 55; 56; 57).

**Figure 1:**
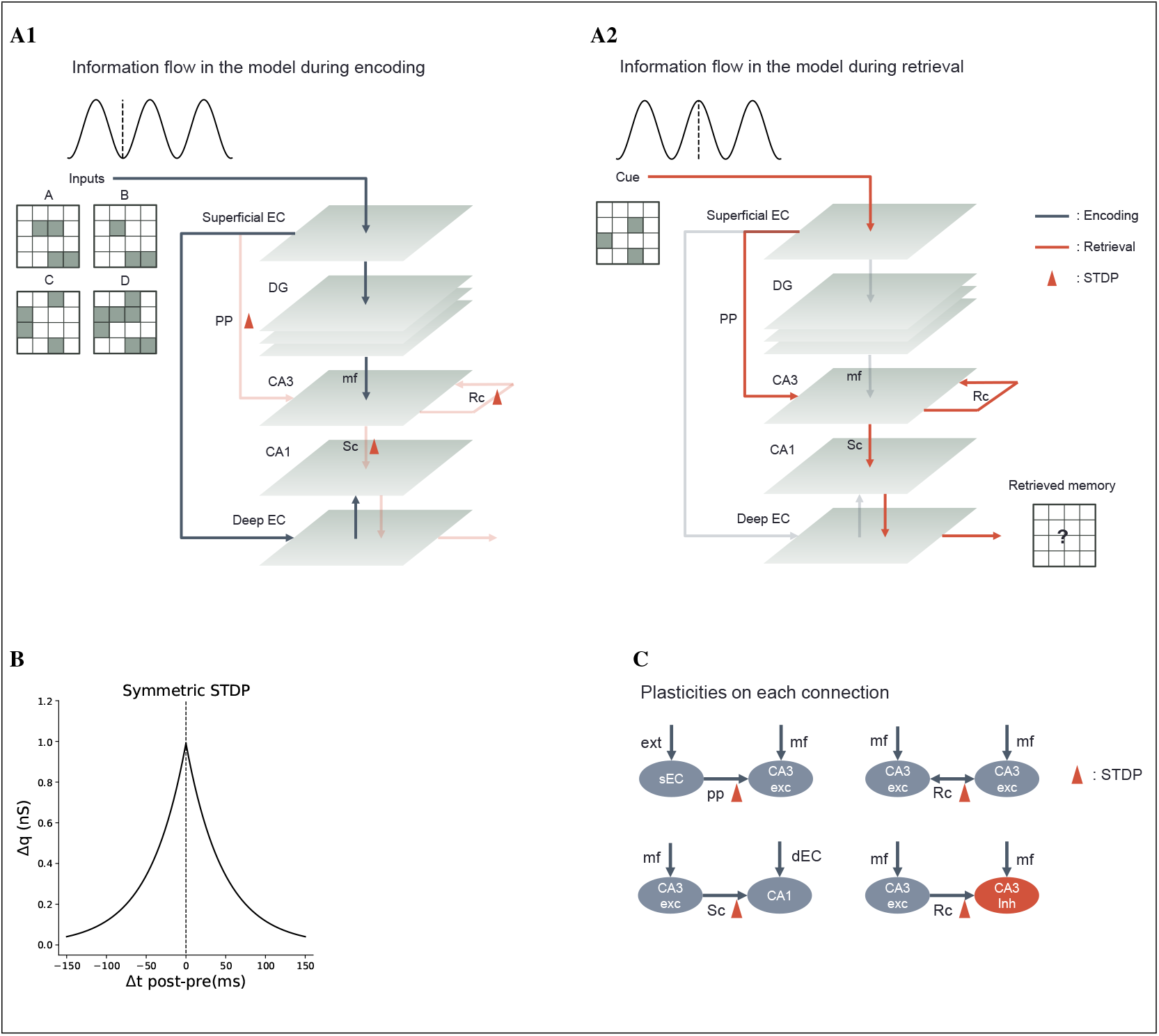
Overview of model architecture and learning. **(A)** The model includes hippocampal subfields: DG, CA3, and CA1, as well as the superficial and deep layers of the EC. The model operates in two distinct phases aligned with theta oscillations, where encoding and retrieval are processed separately. **(A1)** Information flow during the encoding phase. The superficial EC sends inputs to the DG and deep EC, but not directly to CA3. Synaptic plasticity occurs at the silenced connections during encoding, specifically at the PP from the superficial EC to CA3, Rc, and Sc. **(A2)** Information flow during the retrieval phase. The superficial EC sends inputs directly to CA3, which retrieves memories in the deep EC through Rc, Sc, and CA1. **(B)** Symmetric STDP kernel used during the encoding phase. **(C)** Synaptic plasticity across various connections. Abbreviations: ext: external input; mf: mossy fiber; exc: excitatory neuron; inh: inhibitory neuron.

During the retrieval phase, all connections silenced during encoding are reactivated, while those active during encoding are silenced. The superficial EC sends inputs to CA3 via the direct PP, leading to competition among neural assemblies and ultimately the emergence of a dominant engram for pattern completion. During the competition, noise was introduced to CA3 excitatory neurons at a frequency of 3.5 Hz, following a Poisson distribution, which is an essential component for the attractor network (58). This noise simulates the variability and spontaneous activity present in biological neural networks, helping to stabilize the attractor dynamics and prevent the network from getting stuck in suboptimal states. The winning assembly in CA3 then activates CA1, which subsequently triggers neurons in the deep EC completing the retrieval process (Fig 1A2). Each phase lasts 120 ms, allowing sufficient time for both information storage and retrieval. Because there is no interaction between the encoding and retrieval phases in the model, we did not synchronize these phases with oscillatory cycles for most tasks, nor did we base oscillation generation on biological processes such as those in the medial septum (59; 60; 61; 62; 63; 64). Instead, we focused on simplifying the model by emphasizing distinct information flows for each phase.

### 3.2 Engram formation with selective inhibition through heterosynaptic plasticity

This section details the mechanism of engram formation in the model, with a focus on the novel role of selective inhibition induced by heterosynaptic plasticity. Mossy fibers from the DG project to both pyramidal neurons and inhibitory neurons in the CA3 region, facilitating sparse distributed coding (33; 65; 66; 67; 68; 69). These fibers transmit the outputs of pattern separation performed by the DG. We specifically concentrated on mossy fiber inputs to inhibitory neurons in CA3 due to their role in feedforward inhibition, which regulates pyramidal neuron activity (70; 71; 72; 73; 74; 4; 75). Our hypothesis extends beyond the conventional view of feedforward inhibition as a sparsification mechanism (33; 34). We propose that feedforward inhibition, facilitated by mossy fibers, also supports selective inhibition, enhancing competition between neural assemblies through heterosynaptic plasticity.

Heterosynaptic plasticity, wherein synaptic changes at one set of synapses affect another, is well-documented (35; 36; 37; 38). This form of plasticity occurs when the strength of a synapse changes due to activity at a nearby synapse. For example, activating one synapse can strengthen or weaken neighboring synapses, even if they are not directly involved., even if those synapses were not directly activated. This mechanism enables more complex forms of learning compared to traditional Hebbian plasticity. Multiple studies have demonstrated that heterosynaptic plasticity occurs in the direct PP, Rc, and Sc pathways (35; 36; 37; 38).

However, most of the existing evidence pertains to excitatory neurons. In light of this, we hypothesize that heterosynaptic plasticity also occurs at excitatory to inhibitory synapses in CA3. Building on this evidence and our hypothesis, the proposed model incorporates heterosynaptic plasticity at various types of synaptic connections in CA3, including excitatory to excitatory, and excitatory to inhibitory connections (Fig 1C). During the encoding phase, processed outputs from DG stimulate both excitatory and inhibitory neurons in CA3 via mossy fibers. Inhibitory neurons then suppress most of the excitatory neurons, irrespective of their activity. The remaining excitatory neurons form a neural assembly with strengthened recurrent circuits through heterosynaptic plasticity, reinforcing their excitatory connections (Fig 2A). Additionally, these excitatory neurons form associations with activated neurons in superficial EC and CA1 by strengthening direct PP and Sc connections, respectively. The strengthened direct PP provides feature-specific cues from superficial EC to CA3, while the strengthened Sc transmits the reactivated memory trace from CA3 to CA1, completing the retrieval pathway (Fig 1A2).

**Figure 2:**
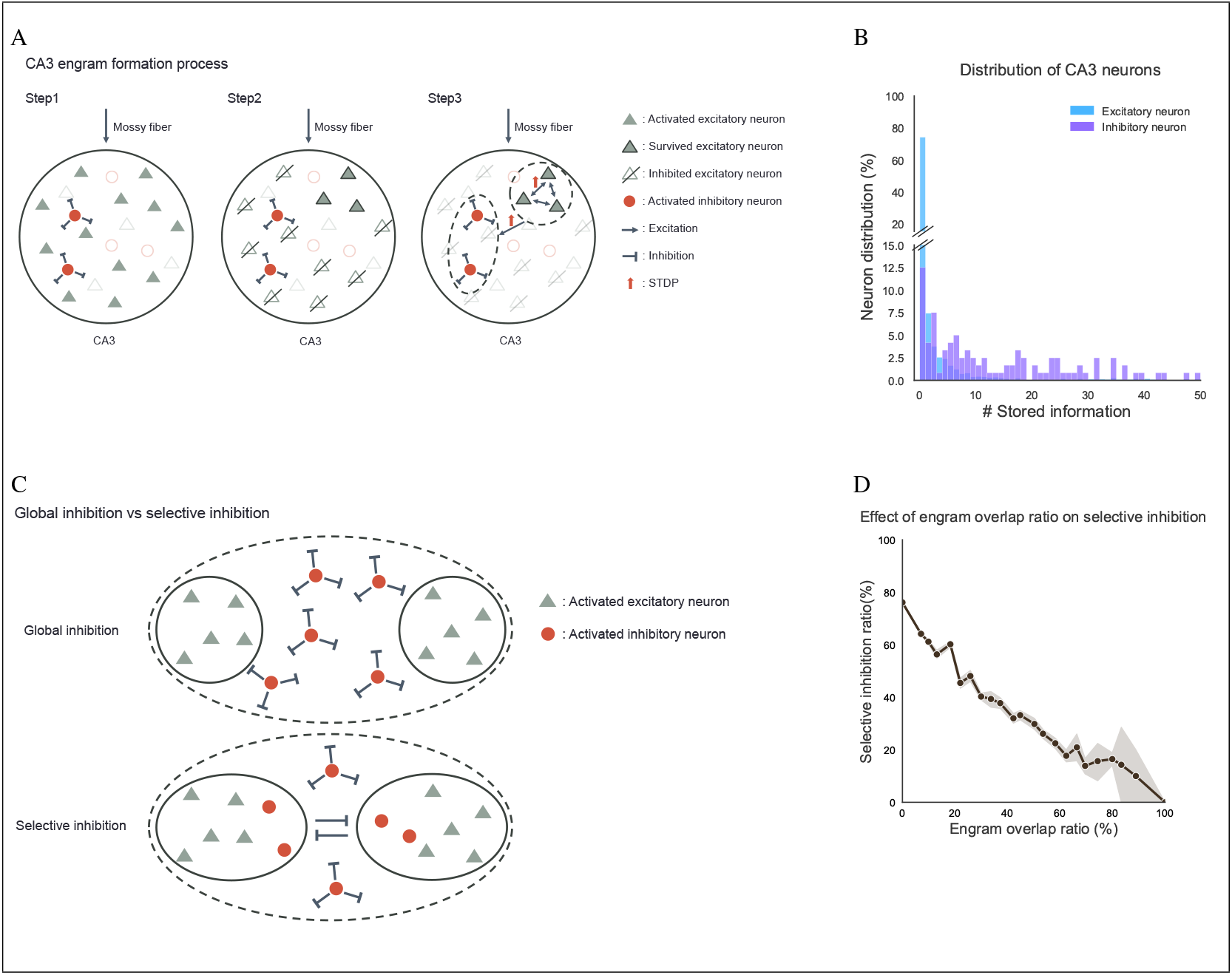
Illustrations and properties for selective inhibition. **(A)** The process of engram formation in CA3 provides the formed engram with the capability for selective inhibition. Inhibitory neurons activated by mossy fibers inhibit most of the excitatory neurons. The surviving excitatory neurons, along with the activated inhibitory neurons, form a neural assembly through STDP. **(B)** Distribution of CA3 neurons based on the number of stored memories. **(C)** Characterization of CA3 connections with inhibitory neurons for global versus selective inhibition. In the case of global inhibition, inhibitory neurons suppress excitatory neurons without regard to which engram they represent. In contrast, for selective inhibition, inhibitory neurons associated with a specific engram do not inhibit excitatory neurons within that same engram. **(D)** The inhibitory influence of one engram on others as a function of the degree of overlap between engrams.

The proposed learning mechanism ensures that inhibitory neurons within an engram do not inhibit excitatory neurons of the same engram (Fig 2C). However, this does not imply that inhibitory neurons are not limited to a single engram; Rather, they exhibit stimulus selectivity. In our model, we refer to this stimulus selectivity as information selectivity, which reflects the ability of neurons to preferentially respond to and store particular aspects of the encoded stimuli. We assessed the information storage capacity of each CA3 neuron by measuring the amount of information retained after encoding 500 arbitrary inputs. As shown in Fig 2B, most excitatory neurons in CA3 (approximately 85%) store only one or two pieces of information. In contrast, inhibitory neurons display a broader range of storage capacities, with some storing significantly more information. These findings suggest that, while inhibitory neurons tend to be information-selective, they are not limited to storing only a single piece of information, consistent with recent findings that they can show selective responses to stimuli (26; 27; 23)

We also investigated how the overlap between CA3 engrams, specifically the shared excitatory neurons within CA3, affects selective inhibition. To evaluate this, we measured the ratio of suppressed excitatory neurons in each neural assembly due to inhibitory neurons associated with competing assemblies. This approach allowed us to assess the extent to which one engram inhibits others. We generated 500 engrams in CA3, each corresponding to an individual input. Engram pairs were selected based on their overlap ratio (‘Materials and Methods’), which was divided into 25 intervals, each representing a 4% range. The inhibition ratio was averaged for each interval. As shown in Fig 2D, the inhibition power of an engram over other engrams decreased as the overlap ratio between engrams increased, indicating that selective inhibition becomes less effective with higher overlap. This observation highlights the importance of sparse distributed coding in CA3 to maintain effective competition between engrams (6; 7; 15; 9; 10; 17).

Strengthened connections from excitatory to inhibitory neurons facilitate earlier activation of inhibitory neurons during retrieval, selectively suppressing most excitatory neurons except those within the same neural assembly in CA3. This selective inhibition helps guide the retrieval cue to a specific memory by promoting the activation of excitatory neurons within the same assembly. The connection weights from excitatory to inhibitory neurons were initialized with nonzero values (‘Materials and Methods’, Fig 5A3), which allowed global inhibition to occur once a sufficient number of neurons in an assembly began firing via recurrent connections, thereby supporting pattern completion.

We propose that selective inhibition provides targeted suppression of competing engrams, ensuring more precise retrieval by reducing interference, while global inhibition, driven by recurrent activity, maintains overall network stability. This dual mechanism could enhance the accuracy of pattern completion and minimize interference between memories with similar content. To evaluate the stability of pattern completion under natural conditions with selective inhibition, we constructed a DG model to generate CA3 engrams with diverse characteristics, such as varying sizes and degrees of overlap. This model enabled a comprehensive assessment of our inhibition mechanism across different scenarios of engram competition. The following section details the construction of the DG model and evaluates its performance in achieving sparse distributed coding through pattern separation.

### 3.3 Feedforward form of the dentate gyrus and its role in instant spatial pattern separation

To endow the dentate gyrus (DG) with spatial pattern separation capabilities, we modeled its structure based on anatomical features while excluding the recurrent circuits, resulting in a feedforward configuration. Numerous modeling studies have demonstrated the effectiveness of the DG in performing pattern separation (76; 77; 78; 40; 79; 80; 81; 82), and it is well-established that a feedforward DG can achieve this function (40). In the proposed model, the DG consists of two distinct layers: the hilus layer and the granule cell layer (GCL) (Fig 3). Although the biological DG contains multiple neuron types, such as hilar interneurons, mossy cells, basket cells, and granule cells (83; 84; 85; 41), we simplified these into two categories—excitatory and inhibitory neurons—using the Izhikevich neuron model (‘Materials and Methods’). Therefore, both the hilus layer and the GCL contain excitatory and inhibitory neurons (Fig 3).

**Figure 3:**
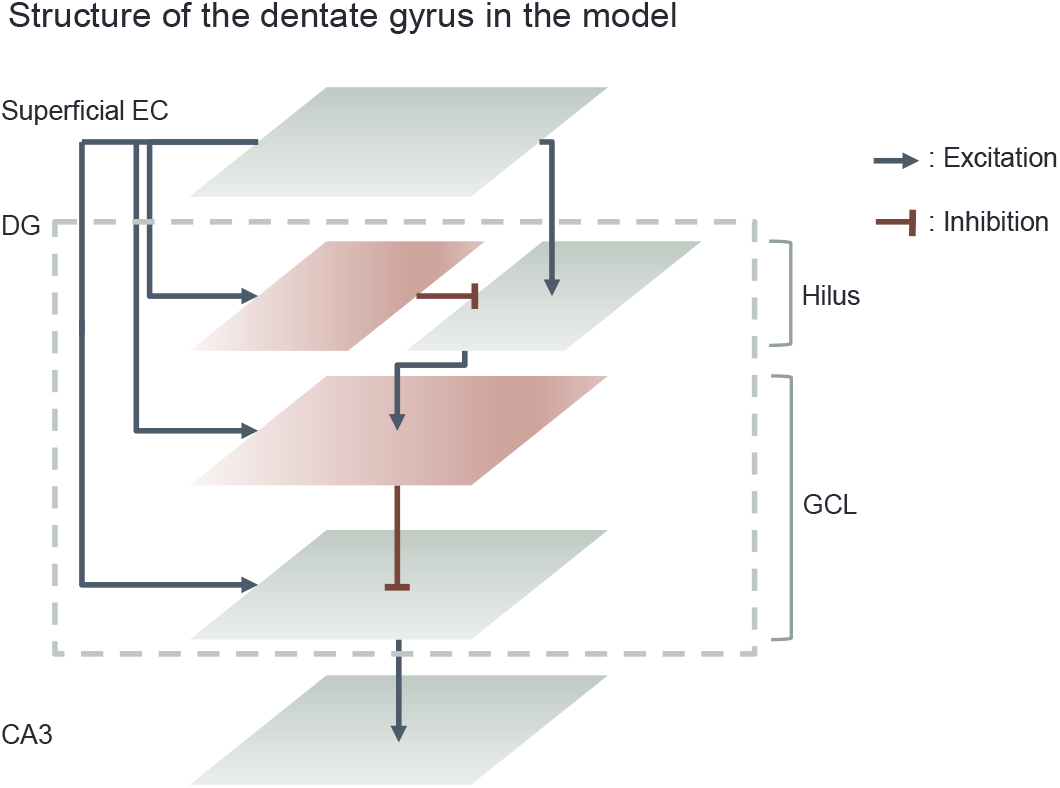
The DG in the model has a feedforward structure consisting of the hilus layer and the GCL. Both layers contain excitatory and inhibitory neurons: excitatory neurons are represented in gray, and inhibitory neurons are represented in red. Superficial EC sends inputs to all types of neurons in the DG. Inhibitory neurons in the GCL also receive inputs from excitatory neurons in the hilus layer, which are regulated by inhibitory neurons in the hilus layer. Finally, excitatory neurons in the GCL, regulated by inhibitory neurons within the GCL, send outputs to CA3 via mossy fibers.

During the encoding phase, inhibitory neurons in the GCL receive two types of excitatory inputs: direct input from the superficial EC and input relayed via the hilus layer. The latter is modulated by inhibitory neurons within the hilus layer, which play a crucial role in pattern separation through disynaptic inhibition (82). Both types of input help detect features of the input pattern, leading to the activation of inhibitory neurons in the GCL for most input patterns (Figs 3 and 4A). Excitatory neurons in the GCL are activated when they simultaneously escape inhibition from GCL inhibitory neurons and receive excitation from the superficial EC. This mechanism transforms low-dimensional input into very sparse, high-dimensional representations. The resulting activation patterns of excitatory neurons in the GCL stimulate both excitatory and inhibitory neurons in CA3 via mossy fibers, creating sparse distributed representations. We passed 10 arbitrary inputs through the DG, and Fig 4A illustrates how these inputs are separated within the excitatory neuron population in the GCL.

**Figure 4:**
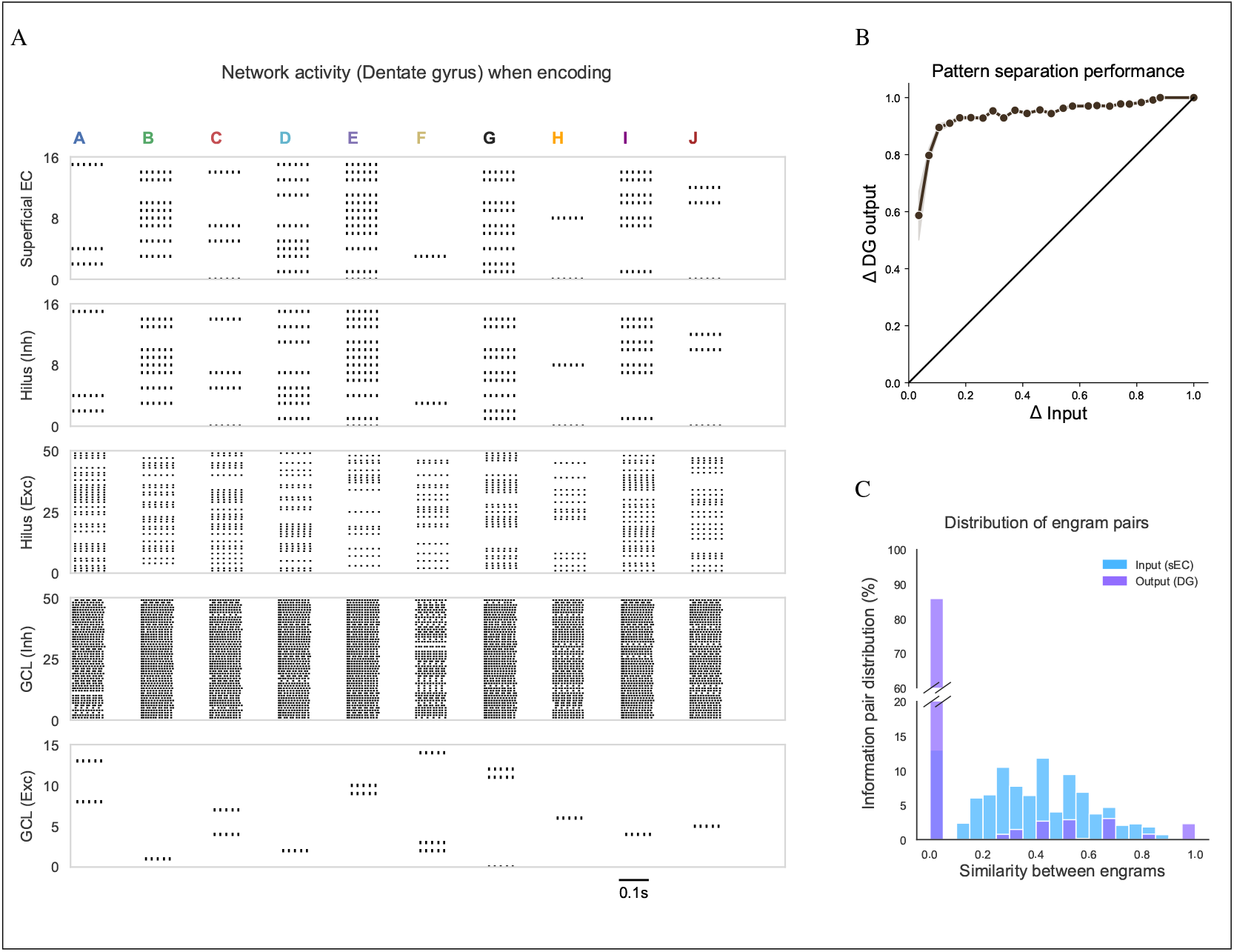
DG spike raster and pattern separation performance. **(A)** Raster plot showing DG activity during the encoding of 10 different inputs (A to J). The activity is displayed for various regions, including the superficial EC, excitatory and inhibitory neurons in the hilus layer, and excitatory and inhibitory neurons in the GCL. For better visualization, 50 neurons are arbitrarily selected from the 100 excitatory neurons in the hilus layer and the 400 inhibitory neurons in the GCL. Additionally, 16 excitatory neurons in the GCL, involved in encoding the 10 inputs (from a total of 800 neurons), are shown. **(B)** Pattern separation performance of the DG. The graph plots the separation performance against input similarity. **(C)** Distribution of engram pairs based on the similarity between activation patterns in the superficial EC (input) and those of excitatory neurons in the GCL of the DG (output).

To evaluate the pattern separation performance of our DG model, we presented 1,000 input patterns and observed the activation patterns of excitatory neurons in the GCL. We calculated the overlap degree for each pair of patterns using the measure suggested by (86), with the discrimination value defined as one minus the overlap value (‘Materials and Methods’). The output discrimination values were averaged within each of the 25 windows, with each window spanning 0.04 units (Fig 4B). This figure shows that the output discrimination values rapidly approach 1.0 when input discrimination values exceed approximately 0.1, indicating effective pattern separation. High output discrimination is maintained for input discrimination values above this threshold. Fig 4C illustrates the distribution of input and output pairs based on their similarity, comparing input pairs from the superficial EC with output pairs from the DG. The analysis indicates that a significant majority of input pairs (over 85%) are fully separated, as reflected by the high percentage of output pairs with minimal similarity. Both Figs 4B and 4C underscore the DG’s spatial pattern separation ability in the model, demonstrating that a feedforward form is sufficient for this function even without any learning.

### 3.4 Sparse distributed coding for memory storage and retrieval

The proposed model, equipped with a feedforward structure of the DG, successfully separates patterns and projects inputs into a high-dimensional space. This process contributes to the sparse distributed representation of memories in CA3, allowing for a diverse range of neural assembly sizes and overlap degrees among assemblies. To assess the storage capacity of CA3, we examined the number of distinct engrams formed relative to the number of inputs (Fig 5C). The results show a consistent increase in the number of engrams, rising by approximately 200 as the number of inputs increases from 600. This indicates that CA3 has sufficient capacity to adapt to increasing input demands without saturating, demonstrating the efficiency of the DG-CA3 network in handling diverse inputs. This capacity of the proposed model is adequate for examining the stability of pattern completion under selective inhibition induced by heterosynaptic plasticity at excitatory to inhibitory connections.

**Figure 5:**
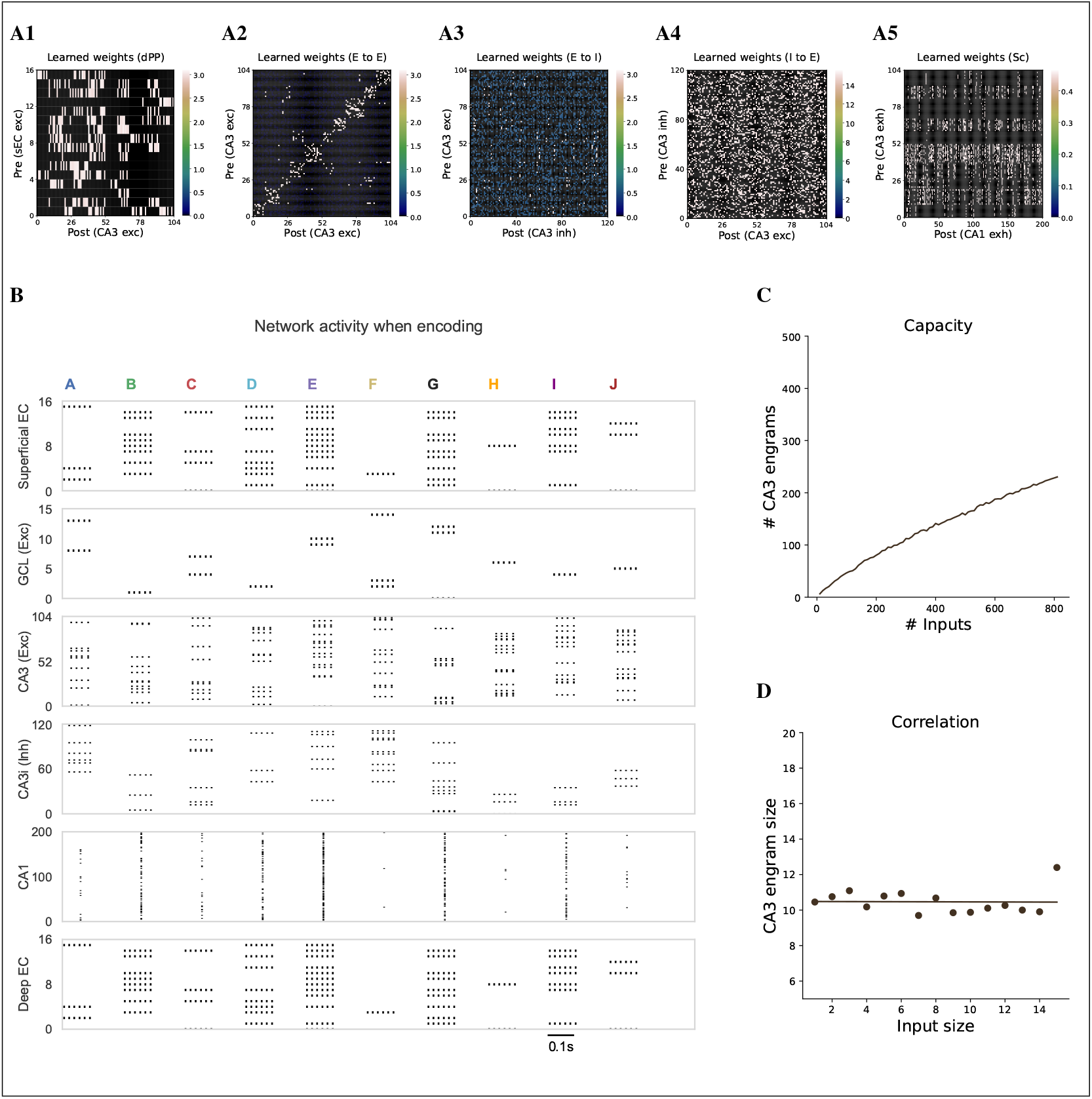
Encoding the various inputs. **(A)** Weight matrices learned through STDP during encoding. Weights represent the fixed connection weights multiplied by the learned peak conductances. Although there are 2,400 CA3 excitatory neurons, only those involved in the encoding of 10 specific examples are selected and arranged for visualization; E, excitatory; I, inhibitory. **(A1)** Learned direct PP weight matrix from superficial EC neurons to CA3 excitatory neurons. **(A2)** Learned excitatory Rc weight matrix. **(A3)** Learned Rc weight matrix from CA3 excitatory neurons to CA3 inhibitory neurons. **(A4)** Learned Rc weight matrix from CA3 inhibitory neurons to CA3 excitatory neurons. **(A5)** Learned Sc weight matrix from CA3 excitatory neurons to CA1 neurons. **(B)** Raster plot showing network activity during the encoding of 10 different inputs (A to J). The activity is depicted for different regions, including the superficial EC, excitatory neurons in the GCL, CA3 excitatory and inhibitory neurons, CA1, and deep EC. Other types of neurons in the DG are omitted for clarity. As in **(A)**, only the CA3 excitatory neurons involved in the 10 specific examples are selected for visualization. **(C)** CA3 capacity is depicted as the number of engrams formed versus the number of inputs. **(D)** Correlation between input size and CA3 engram size, with a linear regression.

Additionally, we analyzed the correlation between the size of the input pattern and the size of the neural assembly in CA3. The size of the patterns in each subregion is defined as the number of firing neurons in the superficial EC and CA3 regions, respectively. Fig 5D shows no correlation between input size and engram size in CA3, indicating that each CA3 engram consistently consists of around 10 excitatory neurons, regardless of input scale. This independence implies that CA3 can efficiently allocate resources without being influenced by input complexity, highlighting the robustness of CA3 engram formation. These results suggest that DG effectively performs pattern separation, enabling CA3 to maintain stable and consistent engrams across varying input conditions.

Figs 5B and 6A illustrate the firing patterns for 10 arbitrary inputs during the encoding and retrieval phases, respectively. Instead of modifying the synaptic weights directly, we updated the peak conductances, which function similarly to conventional synaptic weights (‘Materials and Methods’). This approach keeps the synaptic weights for each connection fixed, thereby maintaining the connectivity ratio. For readability, however, we will refer to the learned weights as the product of the peak conductance and synaptic weights for each connection (Fig 5A).

During encoding, inputs processed by the DG produce distinct firing patterns in excitatory neurons within the GCL, which subsequently activate both excitatory and inhibitory neurons in CA3. With suppression from these inhibitory neurons, the DG output creates sparsely distributed activation patterns among CA3 excitatory neurons (Fig 5A2). In Figure 5A2, most connection weights are concentrated along the diagonal axis, indicating localized connectivity within individual neural assemblies, while others are distributed off-diagonally. This off-diagonal distribution suggests sparsely distributed storage, implying that the learned patterns are not limited to self-connections. In contrast, no diagonal patterns were observed for other connections, particularly those involving inhibitory neurons in CA3 (Figs 5A3 and 5A4). This suggests that while inhibitory neurons are selective to inputs (Fig 2B), they are not specifically constrained to particular engrams.

To observe engram competition during the retrieval phase, we presented an input with 0.5 similarity to both engrams E and G 10 times during retrieval. This input, delivered to the superficial EC, was routed to CA3 via the direct PP, serving as a retrieval cue. Under these conditions, the model successfully retrieved either input G or E in 80% of the trials. Failures were defined as instances where no clear winner emerged from the competition, resulting in interference or no activation in the deep EC (Fig 6A). An analysis of the average firing rate over a 30 ms window for each CA3 neural assembly revealed that only one assembly displayed a significantly high firing rate of approximately 80 to 120 Hz during successful pattern completion, while others exhibited rates of less than about 10 Hz (Figs 6B1 and 6B2). This suggests that the dominant assembly effectively suppressed the activity of competing assemblies. In contrast, during failed pattern completion, no dominant assembly was observed (Fig 6B3).

**Figure 6:**
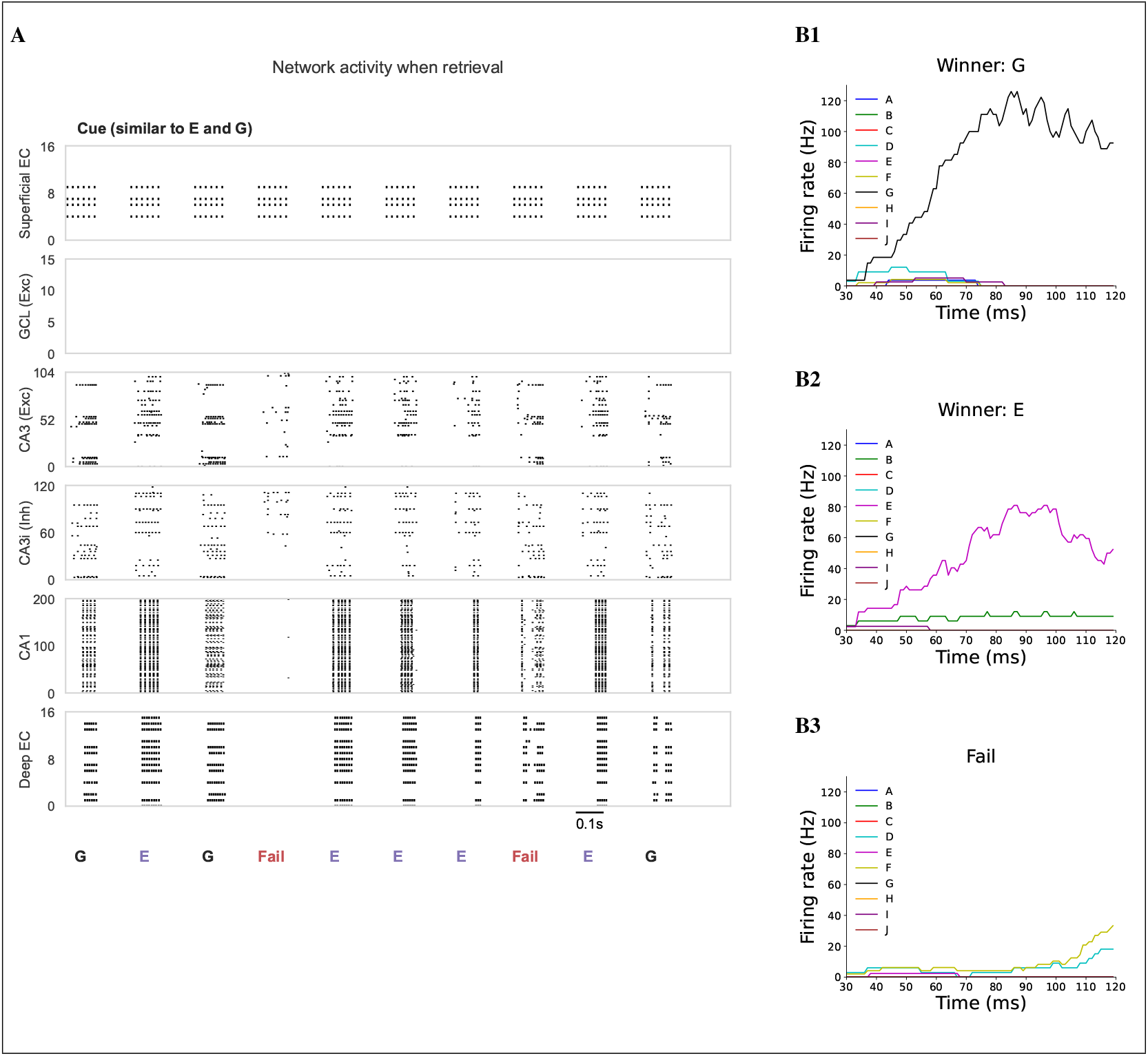
Retrieval for learned inputs. **(A)** Raster plot showing network activity in response to a cue similar to inputs E and G among 10 different inputs (A to J). The depicted layers and neurons correspond to those in Fig 3B. The retrieval cue is presented 10 times, resulting in either successful retrieval (E, G) or retrieval failure, as observed in the activity patterns of the deep EC. **(B)** Firing rate over time for neural assemblies in CA3 during retrieval, illustrating different scenarios of memory retrieval. **(B1)** Example of successful retrieval of input G, showing a dominant firing rate for the neural assembly representing G. **(B2)** Example of successful retrieval of input E, showing a dominant firing rate for the neural assembly representing E. **(B3)** Example of retrieval failure, where no single neural assembly dominates the activity.

These results demonstrate the model’s ability to retrieve memories based on their similarity to retrieval cues. While the DG processes input features to create distinct representations before they reach CA3, the direct PP preserves the original features, allowing for more direct associations with similar memories. To evaluate the function of the direct PP, we deactivated it during retrieval and used mossy fibers to transmit information to CA3. We tested varying percentages of partial cues for each memory to assess the role of the direct PP in capturing input completeness (Fig 7A). Fig 7C illustrates retrieval performance, averaged over 5 samples for each percentage of partial cue. The results show significantly higher success rates when the direct PP is intact, compared to when it is silenced. This difference occurs because the DG perceives the cue as distinct information, activating entirely different neurons in the CA3 region through pattern separation. In contrast, the direct PP stimulates neurons associated with memories similar to the cue (Figs 7B1 and 7B2).

**Figure 7:**
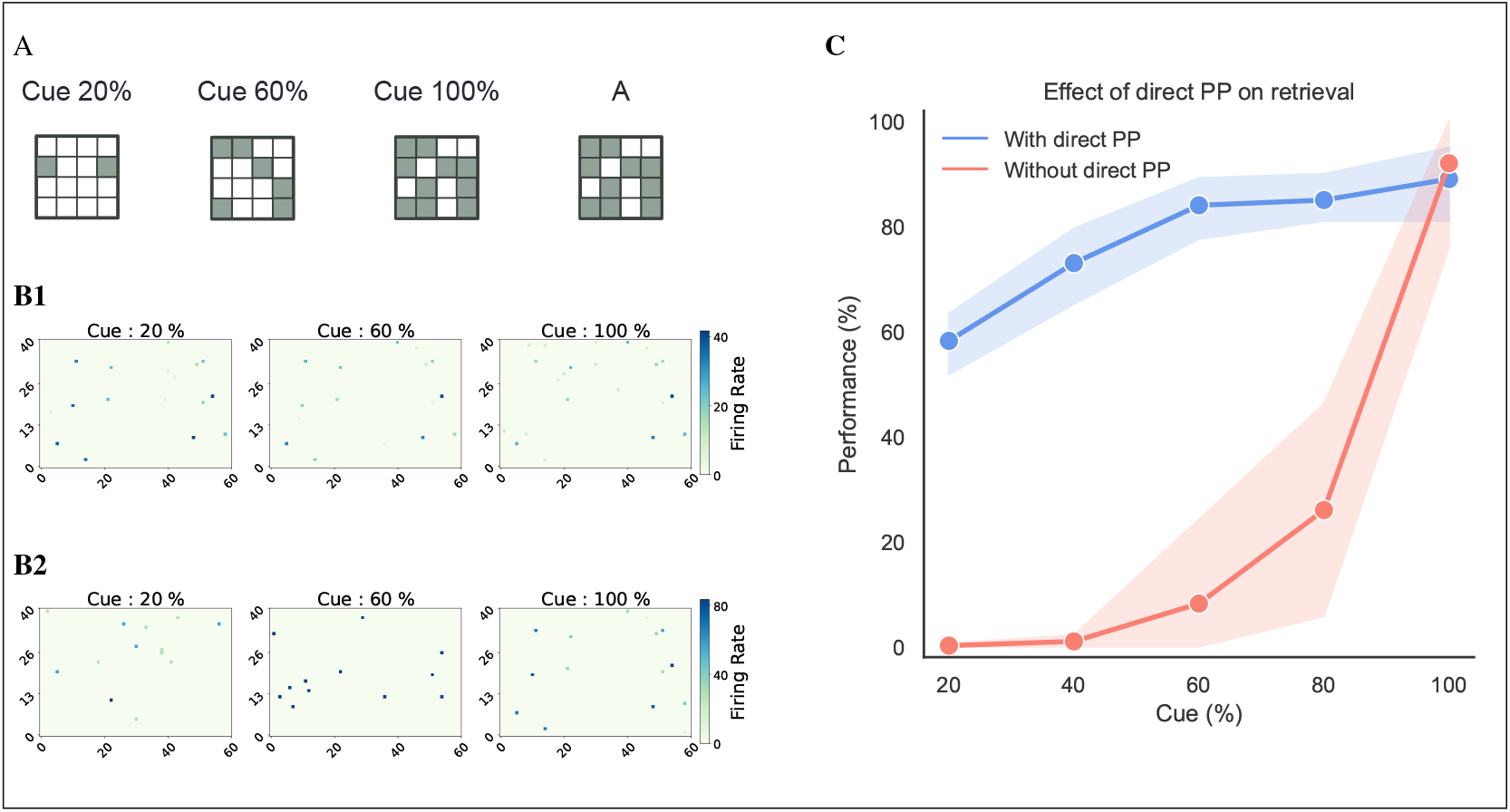
The effect of the direct perforant path for pattern completion during retrieval. **(A)** Illustration of different percentages of partial cues for a learned input A. **(B)** Firing rate heatmap for CA3 neurons in response to a cue during retrieval (120 ms). Since the model may retrieve different outputs in each trial for a given cue, the heatmaps display the firing pattern with the highest likelihood of retrieval across 20 repetitions. **(B1)** Example heatmap for a cue in the intact direct PP condition, showing similar firing patterns regardless of the cue variation. **(B2)** Example heatmap for a cue when the direct PP is silenced, displaying different firing patterns for each cue variation. **(C)** Retrieval performance as a function of the percentage of partial cues under two conditions: with direct PP (blue) and without direct PP (red). Performance is measured as the mean retrieval success rate, averaged over 5 samples for each cue percentage, except for the 100% cue case, which is based on a single sample. The success rate for each sample is averaged over 20 repetitions.

### 3.5 Stability of pattern completion with selective inhibition under various conditions

We demonstrated that the model can encode inputs in a sparse and distributed form through pattern separation by the DG and retrieve these inputs via the direct PP through pattern completion. Furthermore, the independence between CA3 engram size and input size, along with the model’s sufficient capacity for sparse encoding, supports stable pattern separation. The sparsely distributed nature of engrams in CA3 exhibits various properties that may influence competition during pattern completion. We identified four key conditions affecting this competition: (1) disparity in input strength to CA3 engrams, (2) overlap between CA3 engrams, (3) the number of competing engrams for a single cue, and (4) size differences between CA3 engrams, defined by the number of excitatory neurons involved.

To investigate the effects of selective inhibition induced by heterosynaptic plasticity at excitatory to inhibitory connections in CA3, we examined its impact on engram competition and compared retrieval performance with and without selective inhibition. By disabling plasticity at these connections, we removed the mechanism for selective inhibition, making the model reliant solely on global inhibition. This setup allowed a direct comparison between pattern completion with selective inhibition versus global inhibition.

#### 3.5.1 Robust pattern completion against input strength disparity

According to the encoding specificity principle, retrieval is most effective when the retrieval cue closely matches the features present during the initial encoding of the memory (87; 88). However, retrieval can become challenging when competing memories share significant similarities with the retrieval cue, leading to interference. While such interference can naturally result in the generalization of competing memories, it becomes problematic when distinguishing between these memories is necessary. One approach to address this challenge is by providing a biased input to one memory, thereby reducing the similarity of another memory to the cue (89). However, when two memories contain highly similar information, introducing a biased input becomes challenging and may lead to poor retrieval performance. This, in turn, reduces the effectiveness of the dentate gyrus in distinguishing highly similar information.

We explored the role of selective inhibition in stabilizing pattern completion, even when the cue was uniformly similar to the memories encoded by the two competing CA3 engrams. We modulated input bias by adjusting the similarity of cues to one memory relative to another. Specifically, we varied the number of common neurons shared between the cue and the encoded input (Fig 8A). To specifically assess the impact of input bias, we selected 15 pairs of inputs, each stored in CA3 engrams of the same size, with no overlap between engrams (Fig 8B). Each input pair was tested with five different input bias levels, and retrieval was tested 50 times for each level.

**Figure 8:**
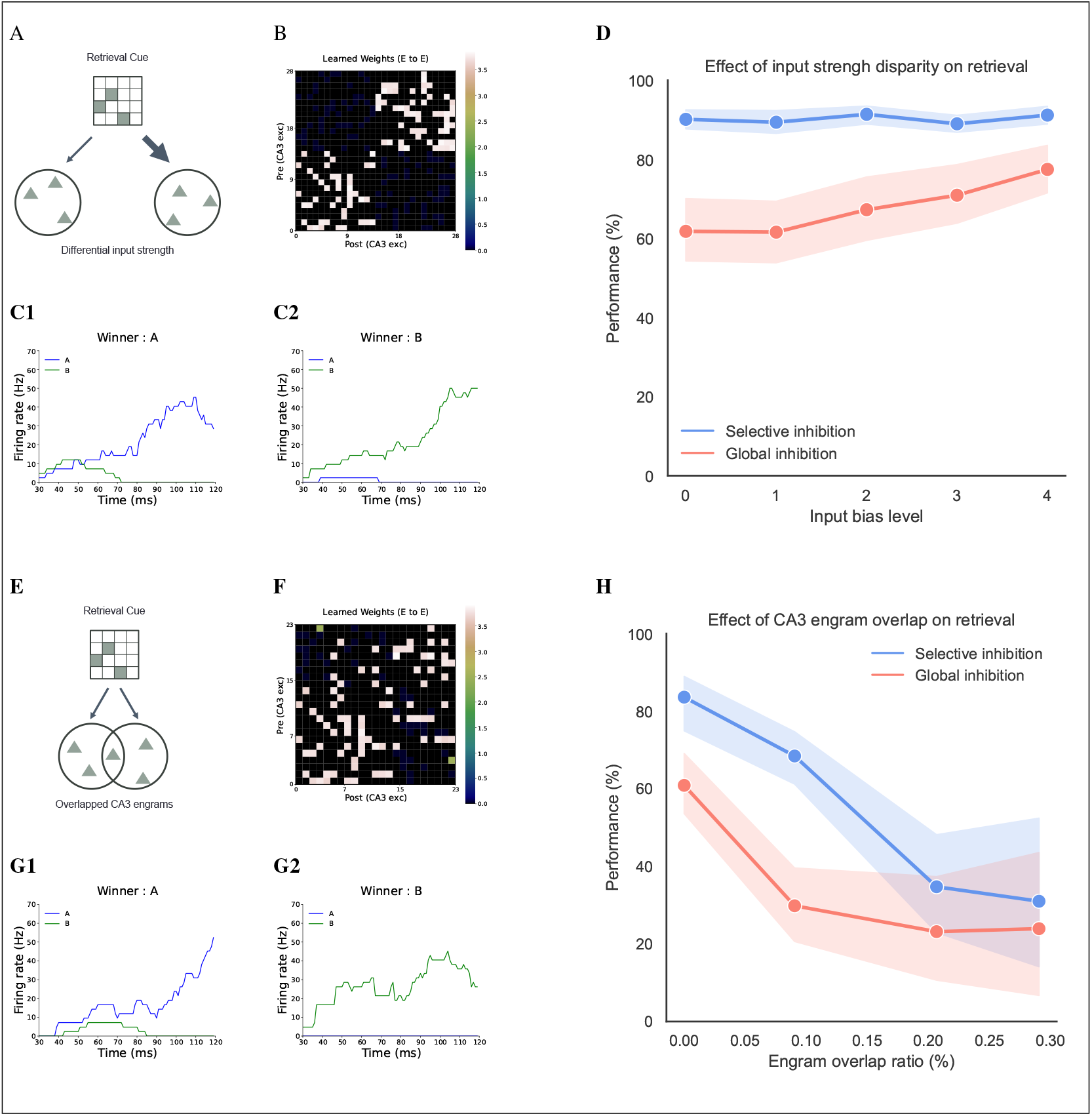
The effect of input strength disparity and overlap ratio between CA3 engrams on retrieval. **(A)** Schematic of the retrieval cue with different input strengths for each engram. **(B)** Example of the learned Rc weight matrix in the input strength disparity task. **(C)** Firing rate over time for neural assemblies in CA3 during retrieval in the input strength disparity task. **(C1)** and **(C2)** show results for the same cue with the same learned samples. **(C1)** Example where engram memory A is retrieved. **(C2)** Example when memory B is retrieved. **(D)** Retrieval performance at different levels of input strength disparity, comparing selective inhibition (blue) and global inhibition (red). Performance is measured as the mean retrieval success rate, averaged over 15 samples for each input bias level. Each sample’s success rate is averaged over 50 repetitions. **(E)** Schematic of the retrieval cue with overlapping engrams. **(F)** Example of the learned Rc weight matrix in the overlap task. **(G)** Firing rate over time for neural assemblies in CA3 during retrieval in the overlap task. **(G1)** and **(G2)** show results for the same cue with the same learned samples. **(G1)** Example when memory A is retrieved. **(G2)** Example when memory B is retrieved. **(H)** Retrieval performance at different overlap ratios between neural assemblies, comparing selective inhibition (blue) and global inhibition (red). Performance is measured as the mean retrieval success rate, averaged over 15 samples for each overlap ratio. Each sample’s success rate is averaged over 50 repetitions.

Our results showed that when the cue was uniformly similar to the memories encoded by two CA3 engrams, retrieval performance under global inhibition was approximately 60%. This suggests that non-biased inputs can hinder engram competition, increasing the likelihood of interference between memories, even when they are fully separated (Fig 8D). In contrast, under selective inhibition, retrieval performance remained consistently high, around 90%, regardless of disparities in input strength This consistency is likely because selective inhibition enhances the contrast between competing memories, effectively amplifying small differences in input strength. These findings indicate that selective inhibition can improve the robustness of memory retrieval in the hippocampus by maintaining pattern completion performance even when cues are not biased toward a particular memory.

#### 3.5.2 Pattern completion with overlapping engrams

Next, we examined the impact of overlap between engrams (Fig 8E). In CA3, engrams are encoded using sparse distributed representations, which can result in varying degrees of overlap. Although there may be mechanisms that could mitigate interference from overlapping engrams, such overlaps still pose a significant risk of interference, potentially reducing retrieval performance. To investigate this effect, we prepared 60 input pairs, each stored in CA3 engrams of the same size, while varying the overlap ratio from 0 to 0.3 (Fig 8F). Each pair of memories received uniformly similar partial cues, and retrieval was tested 50 times.

The results showed that retrieval performance generally declined as the overlap ratio between engrams increased. However, selective inhibition consistently outperformed global inhibition at all overlap ratios (Fig 8H). Specifically, selective inhibition maintained about 70% performance until the overlap ratio reached 0.1, whereas global inhibition drastically decreased to about 30% at the same overlap ratio. Although selective inhibition also experienced a sharp decline, dropping to about 35% as the overlap ratio increased to 0.3, it still performed better than global inhibition. These findings suggest that while the effectiveness of selective inhibition diminishes with higher overlap, it still mitigates interference between overlapping engrams more effectively than global inhibition. This indicates that selective inhibition helps maintain memory integrity, even in the presence of potential interference.

#### 3.5.3 Pattern completion performance relative to the number of competing engrams

We also considered scenarios where multiple memories compete for retrieval from a single cue (Fig 9A). As the number of memories associated with a cue increases, it becomes more challenging for the cue to retrieve specific memories effectively (90; 91), leading to a natural decline in retrieval performance. Nonetheless, studying the effects of selective inhibition in this context is valuable, as it helps us understand its role in managing situations with multiple competing memories. To explore this, we prepared 75 sets of inputs, each stored in CA3 engrams of the same size and without overlap, while varying the number of competing engrams from 2 to 6. (Fig 9B). As in other tasks, each pair of memories received uniformly similar partial cues, and retrieval was tested 50 times.

**Figure 9:**
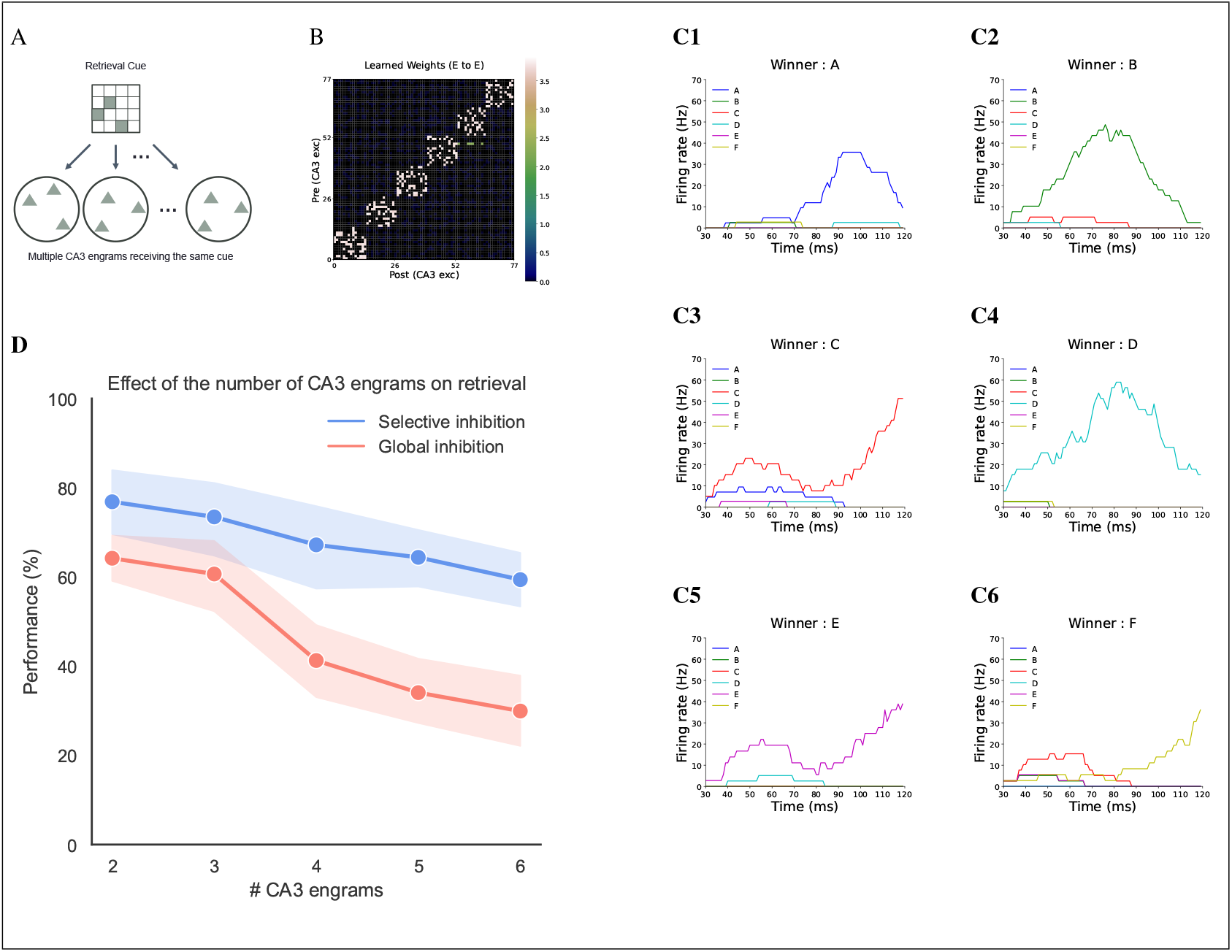
The effect of the number of engrams in CA3 on retrieval. **(A)** Schematic of the retrieval cue with multiple engrams competing for retrieval. **(B)** Example of the learned Rc weight matrix in the task where multiple engrams compete. **(C)** Firing rate over time for neural assemblies in CA3 during retrieval. **(C1)** to **(C6)** show results for the same cue with the same learned samples. **(C1)** Example when memory A is retrieved. **(C2)** Example when memory B is retrieved. **(C3)** Example when memory C is retrieved. **(C4)** Example when memory D is retrieved. **(C5)** Example when memory E is retrieved. **(C6)** Example when memory F is retrieved. **(D)** Retrieval performance across different numbers of engrams, comparing selective inhibition (blue) and global inhibition (red). Performance is measured as the mean retrieval success rate, averaged over 15 samples for each condition involving different numbers of engrams. Each sample’s success rate is averaged over 50 repetitions.

Fig 9D shows that retrieval performance generally decreases as the number of competing engrams increases. For selective inhibition, retrieval performance gradually decreased from about 80% to about 60% as the number of competing engrams increased. In contrast, under global inhibition, performance remained steady at about 60% when the number of competing engrams was 3, but then drastically dropped to about 30% as the number of competing engrams increased to 6. These data suggest that selective inhibition can mitigate some of the negative effects more effectively than global inhibition, even as the number of competing memories increases.

#### 3.5.4 Reduced sensitivity to size variations of the engram in CA3 under selective inhibition

In the model, the size of the engram in CA3 shows no correlation with input information (Fig 5D), suggesting that engram size should not inherently affect pattern completion. Ideally, input similarity should have a greater influence than engram size to prevent scenarios where a less similar memory is easier to retrieve because it just has a larger engram. We investigated the impact of size differences to determine whether a larger engram has an advantage in retrieval and how selective inhibition manages this size disparity compared to global inhibition (Fig 10A). We prepared 75 sets of input pairs, each stored in CA3 engrams with varying sizes and no overlap between engrams (Fig 10B). Each pair of memories received uniformly similar partial cues, and retrieval was tested 50 times.

**Figure 10:**
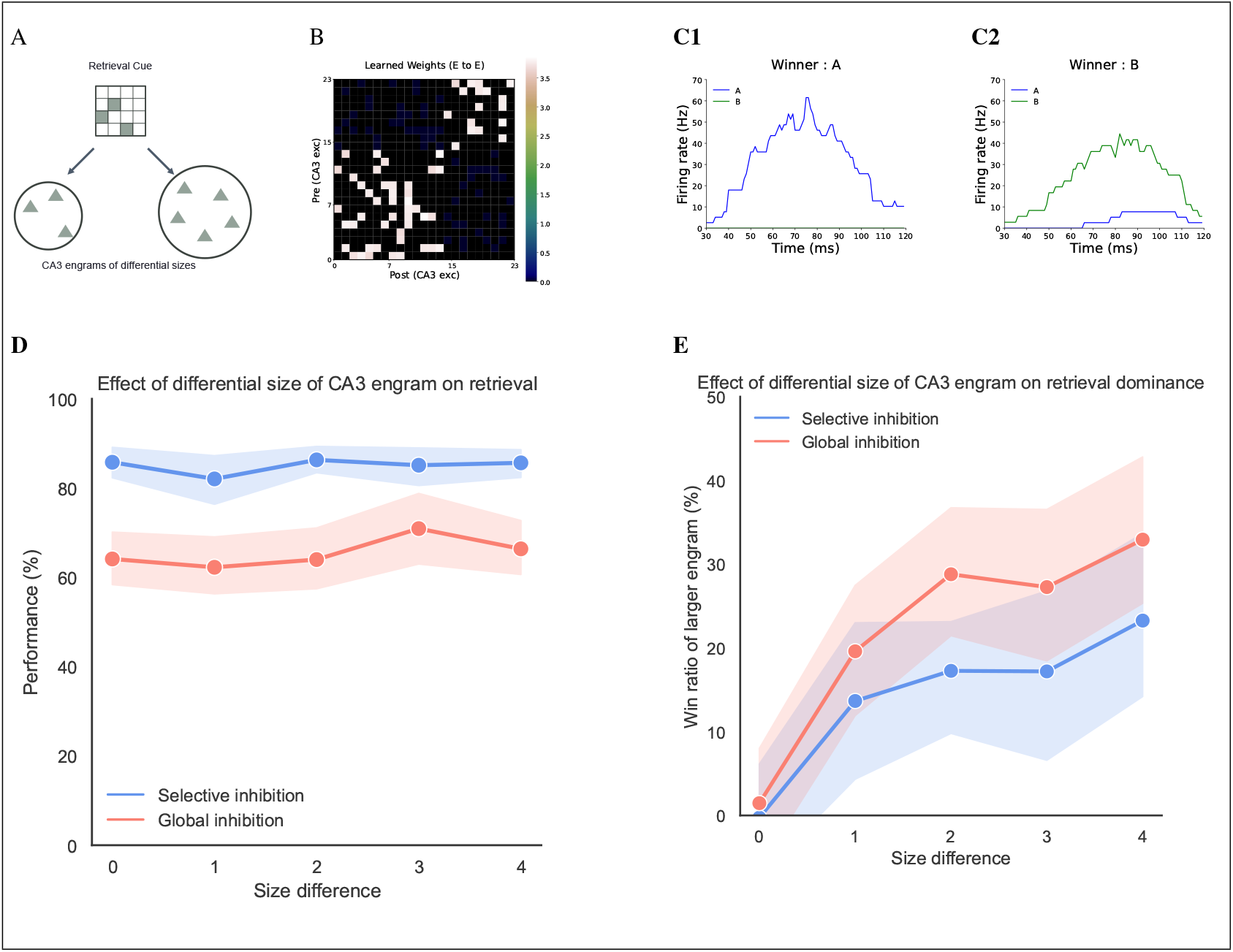
The effect of the size of engrams in CA3 on retrieval. **(A)** Schematic of the retrieval cue with engrams of different sizes. **(B)** Example of the learned Rc weight matrix in the task where engrams have varying sizes. **(C)** Firing rate over time for neural assemblies in CA3 during retrieval. **(C1)** and **(C2)** show results for the same cue with the same learned samples. **(C1)** Example when memory A is retrieved. **(C2)** Example when memory B is retrieved. **(D)** Retrieval performance as a function of engram size differences, comparing selective inhibition (blue) and global inhibition (red). Performance is measured as the mean retrieval success rate, averaged over 15 samples for each size difference. Each sample’s success rate is averaged over 50 repetitions. **(E)** Retrieval dominance of memories stored in larger engrams across different size differences, comparing selective inhibition (blue) and global inhibition (red). Retrieval dominance indicates the percentage of times the memory stored in the larger engram is retrieved.

The results show that selective inhibition consistently maintained about 80% retrieval performance, outperforming global inhibition, which achieved about 60% (Fig 10D). This result is consistent with the findings in Fig 8D. However, the size of the engram influenced which memory was retrieved. We conducted 50 trials to investigate which memories were retrieved and calculated the retrieval ratio by comparing the difference in retrieval counts for each memory type to the total number of successful retrievals. Memories stored in larger engrams were more likely to dominate the retrieval process. This effect was particularly pronounced under global inhibition, with larger engrams accounting for about 30% of retrievals, compared to about 20% under selective inhibition (Fig 10E). This competition revealed a tendency for memories stored in larger engrams to be retrieved more frequently, highlighting the potential limitations of global inhibition in managing engram competition.

Our findings demonstrate that selective inhibition provides a robust mechanism for pattern completion across various conditions that affect engram competition. It enhances retrieval performance regardless of input strength disparity, mitigates the negative effects of overlap between engrams, maintains higher performance levels as the number of competing memories increases, and effectively manages size differences between engrams. These results underscore the functional advantages of selective inhibition over global inhibition in reliable memory retrieval. They support our hypothesis that connections from excitatory to inhibitory neurons can be learned during encoding via heterosynaptic plasticity and suggest a potential role for this mechanism in managing complex memory dynamics within the hippocampus.

### 3.6 Neuronal representations across hippocampal subregions during pattern separation and pattern completion task

We expanded our investigation to focus on pattern separation across the entire hippocampal model, building on our observations of stable pattern completion under various conditions facilitated by selective inhibition. Previous studies have shown that distinct hippocampal subregions exhibit different patterns of neuronal activation (13; 10). To explore this, we conducted 105 trials, varying the similarity between inputs A and B (Figs 11A and 11B). Input A was presented during a single encoding phase lasting 120 ms, while input B was presented over 7.2 seconds across 30 phases, encompassing both encoding and retrieval (Fig 11A).

**Figure 11:**
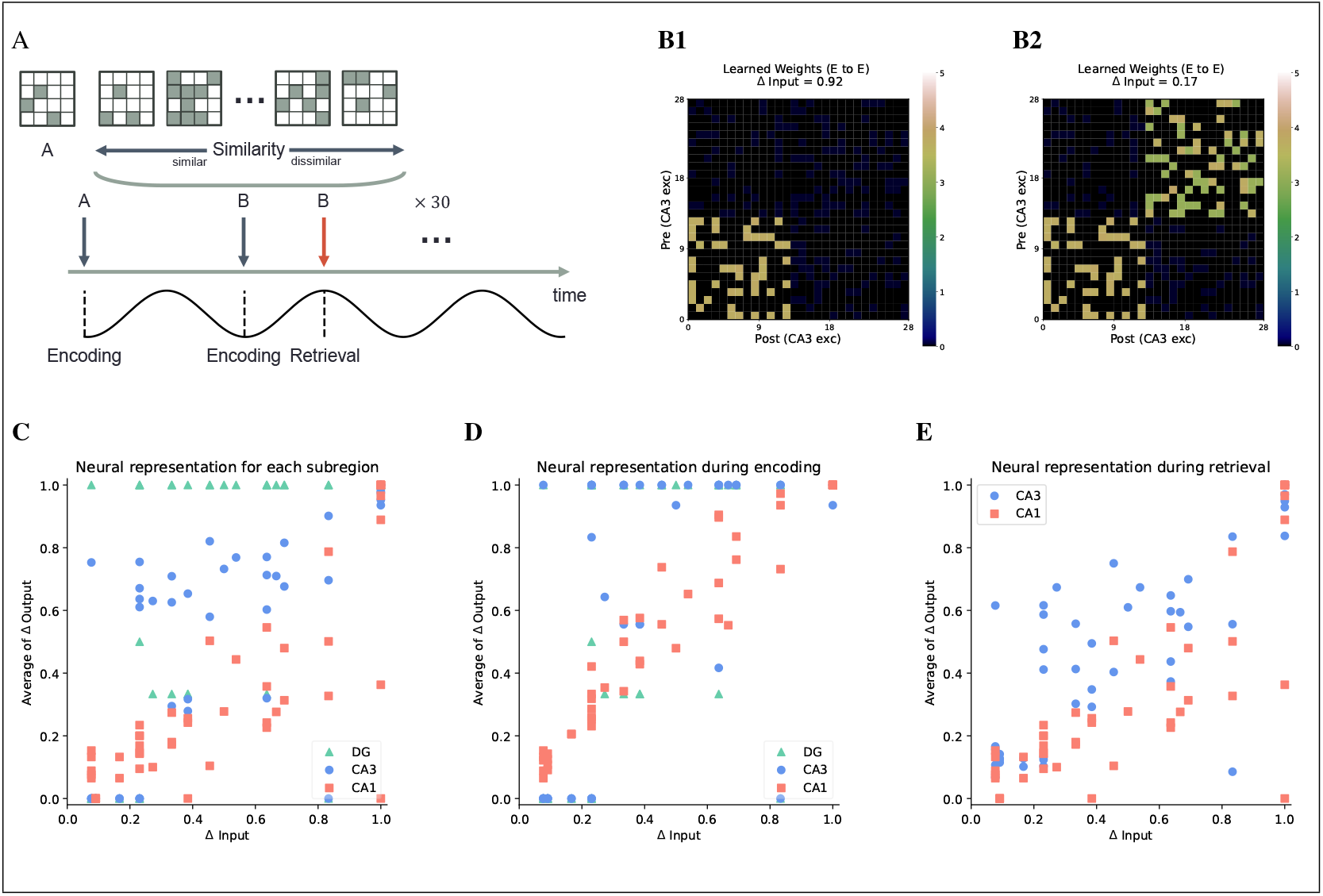
**(A)** Schematic of the pattern separation task. Input A is presented to the model during a single encoding phase (120 ms), while Input B is presented over 7.2 seconds (30 phases), encompassing both encoding and retrieval phases. Input B’s similarity to Input A is varied across 105 trials to evaluate the model’s pattern separation ability. **(B)** Examples of learned Rc weight matrices for **(B1)** highly similar (Δ Input = 0.92) and **(B2)** less similar (Δ Input = 0.17) input patterns. **(C)** Neural representation changes across subregions in the model as a function of input discrimination (Δ Input). The Δ outputs are averaged over 30 phases. **(D)** Neural representation changes across subregions in the model as a function of input discrimination (Δ Input) during the encoding phase. **(E)** Neural representation changes across subregions in the model as a function of input discrimination (Δ Input) during the retrieval phase.

To quantify the discrimination between neuronal patterns A and B, we averaged the discrimination values across 30 phases. Our analysis revealed distinct computational characteristics for each hippocampal subregion. The DG exhibited robust pattern separation, effectively distinguishing input patterns, particularly when input dissimilarity (Δ input) exceeded about 0.2. In contrast, CA1 showed a linear relationship between input and output discrimination, indicating a more gradual differentiation process (Fig 11C). This finding aligns with evidence suggesting that the DG has a unique capacity to discriminate between similar inputs (92; 8). In the CA3 region, Δ output tended to be lower than in CA1 when Δ input was below about 0.2, indicating reduced efficiency in pattern separation for highly similar inputs. However, as Δ input increased beyond 0.2, CA3’s Δ output surpassed that of CA1, suggesting enhanced separation capabilities for more distinct inputs (Fig 11C). These results are consistent with previous studies (16; 93; 94; 13; 10). We investigated these Δ output patterns by separating the encoding and retrieval phases to understand the reasons for these behaviors. During encoding, CA3 closely mirrored the pattern observed in the DG, while CA1 displayed a more linear response (Fig 11D). This similarity arises because CA3 receives input from the DG during the encoding phase in the model. At Δ input levels below approximately 0.2, the reduced output discrimination in CA3 is attributed to a failure of pattern separation in the DG, where the model incorrectly perceived two distinct inputs as identical. This finding explains the lower output discrimination in CA3 compared to CA1, as shown in Fig 11C.

During retrieval, the pattern in CA3 more closely resembled that of CA1 than it did during encoding, consistent with the model’s design, where CA3 transmits information to CA1 (Fig 11E). Notably, the Δ output of CA3 during retrieval was lower than during encoding, despite input B being provided as an intact cue. This resulted in CA3 appearing to have lower pattern separation performance than the DG in Fig 11C, even though CA3 retained the outcomes of the DG’s pattern separation during encoding (Fig 11D). This effect was observed even at high Δ outputs (around 0.8), where input B did not consistently converge to its own representation due to the competing influence of pattern completion involving input A (Fig 11E).

To gain deeper insights into the dynamics of pattern completion, we designed a specific task to assess this process. We investigated how varying degrees of similarity between retrieval cues and stored memories impact retrieval accuracy. Specifically, we used an input pair with zero similarity (A and B) as the stored memories. During retrieval, we introduced cues with graded similarities to inputs A and B, maintaining a linear proportional relationship between each cue’s similarity to A and B (Fig 12A). For example, if a cue was 70% similar to A, it would be 30% similar to B. This linear progression enables a more precise analysis of the transition between retrieving different memories compared to arbitrary cue selection.

**Figure 12:**
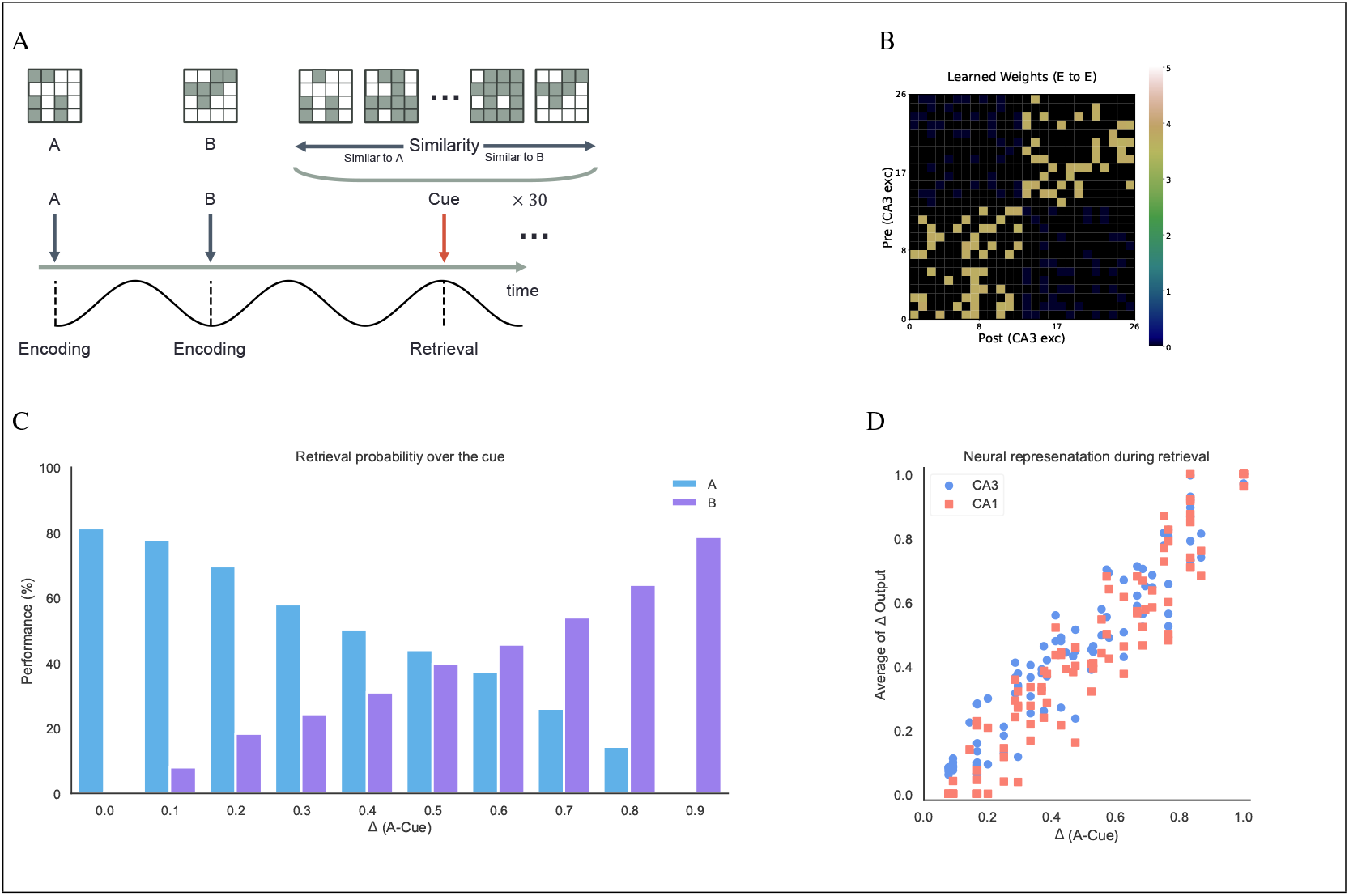
**(A)** Schematic of the pattern completion task. During the encoding phase (120 ms), inputs A and B are presented to the model with a similarity of 0. In the retrieval phase, 100 cues are presented, each having a linearly proportional relationship in their similarity to A and B, ensuring that the sum of these similarities equals 1. Each cue is presented for 7.2 seconds, and 30 retrieval phases are observed without further learning. **(B)** The learned Rc weight matrix after training with inputs A and B. **(C)** Retrieval performance as a function of the similarity between input A and the cue (Δ −A Cue). Performance is measured by the probability of retrieving Input A (blue) or Input B (purple) in response to the presented cue across 30 retrieval phases. **(D)** Neural representation changes relative to the discrimination between input A and the cues (Δ A−Cue) during retrieval. The Δ outputs are averaged over 30 phases.

The results demonstrated a strong linear relationship between cue similarity and retrieval accuracy. Given that inputs A and B were entirely dissimilar, both CA3 and CA1 exhibited highly similar linear response patterns. Notably, the cues did not converge on a dominant memory, even when there was a small degree of similarity to one of the inputs. This finding suggests that the hippocampal model’s pattern completion mechanism is linearly influenced by the degree of similarity between the retrieval cue and the stored memory (Figs 12C and 12D). The linear relationship indicates that even minimally similar cues can lead to accurate memory retrieval, highlighting the model’s sensitivity and precision in pattern completion. These insights into the specific mechanisms of pattern completion within our hippocampal model deepen our understanding of memory processes.

## 4 Discussion

We developed a spiking neural network model based on theta oscillation theory that captures the structural and connectivity properties of the hippocampus. In our exploration of memory processing with this model, we focused on the contributions of inhibitory neurons in CA3. Our findings suggest that the activation of these neurons, along with heterosynaptic plasticity during encoding, can lead to selective inhibition during retrieval. Thus, the role of inhibitory neurons in CA3 includes both selective inhibition and sparse distributed coding. In the proposed model, inhibitory neurons associated with specific neural assemblies in CA3 selectively suppress excitatory neurons involved in competing assemblies.

Interestingly, inhibitory neurons do not suppress excitatory neurons within the same engram. This occurs because only excitatory neurons that are not suppressed by inhibitory neurons survive to form neural assemblies. These roles of inhibitory neurons in memory processing are supported by experimental evidence, including (1) feedforward inhibition of the dentate gyrus via mossy fibers, (2) plasticity within inhibitory neurons, (3) heterosynaptic plasticity across the hippocampus, and (4) the segregation of encoding and retrieval phases by theta oscillation.

### 4.1 Model validation for memory formation

During encoding in the model, inhibitory neurons in CA3 receive inputs from the DG via mossy fibers. These inhibitory neurons then suppress most excitatory neurons in CA3, determining which neurons remain active. Some observations from the literature, although not all fully specific to the hippocampus, align with the proposed model: Granule cells in the DG are known to recruit feedforward inhibition via mossy fibers (70; 73; 74). Parvalbumin-positive (PV+) inhibitory neurons in the hippocampus are crucial for memory storage (31; 32; 22). Inhibitory neurons influence the size of the engram (71; 72; 4; 75). These experimental findings underscore the importance of inhibitory neurons in memory formation, indicating that the model effectively captures their functional and structural roles.

In the model, inhibitory neurons have stimulus selectivity because the DG processes and separates external inputs before transmitting them to CA3. This feature parallels the stimulus selectivity observed primarily in inhibitory interneurons within the visual and auditory cortices (26; 27; 23), implying that similar mechanisms may exist in the hippocampus. In this context, inhibitory neurons in sensory areas play a more active role in information processing than previously thought. In line with these findings, the model supports the view that inhibitory neurons in the hippocampus actively participate in both encoding and retrieval processes while exhibiting stimulus selectivity.

Observations of plasticity in inhibitory neurons indicate their synaptic interactions with excitatory neurons (95; 31; 28; 29; 30; 32). In the model, this plasticity enables inhibitory neurons to modulate their synaptic strengths with excitatory neurons, thereby facilitating selective inhibition during engram competition. Selective inhibition positions inhibitory neurons as active participants in retrieval processes, underscoring their potential contribution to pattern completion. To effectively harness the properties of inhibitory neurons mentioned above, the model employs two key mechanisms: theta oscillation theory (43; 49; 50; 51; 52; 53) and heterosynaptic plasticity (35; 36; 37; 38). Theta oscillation theory posits that distinct phases of the theta rhythm modulate different information flows in the hippocampus, with encoding occurring at the trough and retrieval at the peak. The hippocampus maintains these distinct phases through phasic changes in membrane potential (43; 96).

In the model, reflecting these phenomena, synaptic activity related to retrieval is suppressed during encoding. During engram formation in CA3, heterosynaptic plasticity strengthens connections between inhibitory and excitatory neurons, even in the absence of direct stimulation. These mechanisms enable inhibitory neurons to play an active role in both sparse distributed coding for pattern separation and selective inhibition for pattern completion.

Proposed model illustrates the rapid process of hippocampal memory formation, completed in just 120 milliseconds. This rapid storage capability reinforces the model’s functional plausibility by effectively replicating the rapid memory formation observed in the hippocampus (97; 98; 99). Moreover, the model demonstrated stable pattern completion across various realistic retrieval scenarios in the hippocampus, including those with evenly distributed input cues, overlapping engrams, multiple competing engrams, and engrams of varying sizes. This stability suggests that the model effectively captures key aspects of how the hippocampus retrieves memories under different engram configurations.

### 4.2 Alignment with cognitive evidence

The enhanced stability of the model, driven by selective inhibition, allowed us to investigate firing patterns across hippocampal subregions during pattern separation and completion by quantitatively controlling input similarity. The distinct functions of hippocampal subregions during pattern separation and completion are well-documented in the literature (13; 10). The DG is particularly robust in supporting pattern separation (11; 14; 92; 8). In CA1, output discrimination is thought to be linearly related to input discrimination, meaning that output differences increase at a constant rate as input differences increase (94; 13; 10). CA3, however, exhibits more complex dynamics because it participates in both pattern separation and pattern completion. Some studies suggest that CA3 is more involved in pattern separation, while others emphasize its greater role in pattern completion. (93; 16; 94).

The model successfully replicates the distinct roles of hippocampal subregions, demonstrating strong pattern separation in the DG and a consistent linear response in CA1. In CA3, the model aligns with the evidence discussed earlier, displaying either pattern separation or pattern completion depending on the degree of input similarity. We were able to effectively investigate CA3’s behavior by clearly separating the encoding and retrieval phases. When the difference between inputs is small, the difference between output pairs in CA3 is reduced compared to CA1. This occurs because pattern separation in the DG does not reach the discrimination threshold during encoding. As a result, the model interprets the two similar inputs as identical and performs pattern completion.

When the difference between inputs is large, CA3 shows less pattern separation than the DG, but still more than CA1. This reduced separation is likely due to mixed outcomes, where two completely different memories are retrieved. In other words, a cue might still trigger pattern completion for less similar memories, even when it closely resembles a specific memory. This is supported by the observation that retrieval probability depends linearly on the similarity between the cue and the memory. Such linearity prevents the dominance of biased memories during retrieval, partially capturing the complexity of real-world information. Because the proposed model aligns with established experimental data, it may serve as a valuable tool for understanding hippocampal functions. By examining phase-specific dynamics in the model, we gain new insights into how the hippocampus handles conflicting information.

### 4.3 Limitations and future works

One of the main assumptions of our model is that heterosynaptic plasticity in the connections from excitatory to inhibitory neurons in CA3 drives selective inhibition. While our model suggests that this plasticity could be pivotal for selective inhibition, experimental evidence for heterosynaptic plasticity at these specific connections has yet to be demonstrated. Therefore, further experimental investigations are necessary to validate this mechanism.

By simplifying the DG to focus solely on spatial pattern separation with a fixed input activation frequency, we overlook its established roles in temporal and rate pattern separation (100; 79; 86). As a result of this simplification, certain details, such as the number of neurons in each subregion, connection probabilities between subregions, and synaptic weights, do not fully align with biological specifics. Since these differences may impact the memory retrieval process, the observed linear relationship between similarity and retrieval probability in the pattern completion task should be interpreted with caution, warranting further experimental validation and larger-scale modeling.

We limited CA1’s role to reinstatement of original information from CA3 engrams, which overlooks the neuronal diversity and complex functions of CA1 (101; 102). The model disregards cellular-level processes, feedback connections, and molecular signaling pathways, such as those mediated by cAMP response element-binding protein (CREB) (95; 4; 103). Since these elements are key components of memory processes, their omission may limit the quantitative accuracy of the model’s outcomes.

In future work, we will extend the model to incorporate sequential learning, exploring how inhibitory neurons contribute to engram formation and competition across sequences of information. This extension could deepen our understanding of hippocampal function in processing sequences, offering new insights into cognitive processes such as replay, sequence compression, and predictive coding.

### 4.4 Conclusion

Our research highlights the active and dynamic role of inhibitory neurons in CA3 engram formation and stable pattern completion. This work provides a fresh perspective on established hippocampal functions and opens new avenues for experimental investigation. As our understanding of hippocampal circuitry and function continues to evolve, models like ours will be essential in bridging the gap between cellular mechanisms and cognitive phenomena, advancing research in both neuroscience and artificial intelligence.

## 5 Methods and Materials

### 5.1 Model structure

We constructed a spiking neural network model that includes the EC and hippocampal subregions-DG, CA3, and CA1-focusing on the trisynaptic circuit, which connects superficial EC to CA1 via DG and CA3. The model emphasizes the dynamics of memory formation and competition in CA3 while simplifying the monosynaptic pathway between EC and CA1. Key pathways, including mossy fiber projections to CA3 from DG and the direct PP from superficial EC to CA3, are retained. The DG is composed of the hilus and GCL, each containing excitatory and inhibitory neurons, similar to CA3. Detailed parameters can be found in Table 2 and Table 3. We concentrated on spatial pattern processing, excluding temporal sequences to maintain simplicity. The input layer (superficial EC) consists of 16 neurons, each firing at a constant rate of 50 Hz. This configuration allows for 2^16^ unique spatial patterns, providing sufficient variability to assess the model’s performance while maintaining computational feasibility.

Given the focus on spatial patterns, we considered only spatial pattern separation in DG, excluding temporal and rate-based separation mechanisms. This simplification inevitably results in deviations from biologically realistic parameter settings across the broader hippocampal model. These deviations include differences in neuron population sizes, connection probabilities, and synaptic weights within hippocampal subregions (Table 2 and Table 3). Initial parameters were derived from the study by (104) and subsequently optimized through iterative manual adjustments to ensure proper model functioning. Connection weights were initialized with specific values and fixed to preserve connection probabilities during learning. Instead of adjusting these weights directly, we modulated the peak conductance, which influences synaptic efficacy (detailed explanation in the next section).

The CA1 region in the model is simplified to include only excitatory neurons (Table 2). Connections between CA1 and deep EC were designed so that each CA1 neuron maintains consistent bidirectional connections with specific deep EC neurons, based on a defined connection probability (Table 3). Additionally, deep EC receives inputs from superficial EC via a one-to-one connection, providing input information to CA1 during encoding. While this simplification omits some intricate microcircuitry in biological CA1 and its connections with EC (101; 102), it retains the essential function of interfacing between CA3 and deep EC, allowing CA3 engrams to effectively encode and decode information between hippocampal and cortical representations.

We implemented the model using Python with NumPy. The following sections detail the specific neuron models, learning rules, and analysis methods used to investigate memory dynamics within this structure.

### 5.2 Single neuron models

Each neuron was described using the Izhikevich model (42) following as

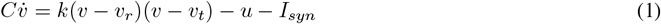

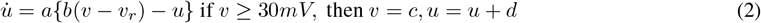

where *C* is the membrane capacitance, *v* is the membrane potential, *v*_*r*_ is the resting potential, *v*_*t*_ is the threshold potential, and *u* is the recovery variable (the difference of all inward and outward voltage-gated currents) (42). *I*_*syn*_ is the summation of synaptic currents from other neurons, detailed in the next section. Each parameter, including the dimensionless values *k, a, b, c*, and *d*, follows the values specified by (42) for different neuron types. Based on this paper, the excitatory neurons use the parameters for pyramidal cells, and the inhibitory neurons use those for basket interneurons (Table 1).

**Table 1:**
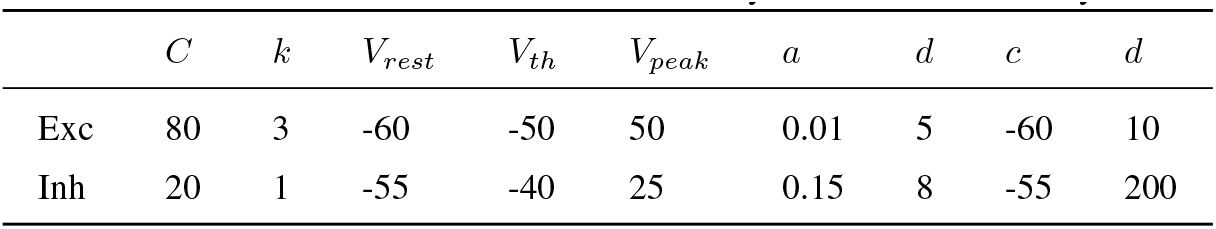
Parameters of Izhikevich model for excitatory neurons and inhibitory neurons.

| | $C$ | $k$ | $V_{rest}$ | $V_{th}$ | $V_{peak}$ | $a$ | $d$ | $c$ | $d$ |
| --- | --- | --- | --- | --- | --- | --- | --- | --- | --- |
| Exc | 80 | 3 | -60 | -50 | 50 | 0.01 | 5 | -60 | 10 |
| Inh | 20 | 1 | -55 | -40 | 25 | 0.15 | 8 | -55 | 200 |

### 5.3 Synapse models

The total synaptic current model also follows (42; 104), which consists of four basic conductances from AMPA, NMDA, GABA_A_, and GABA_B_ channels with the simplified voltage-gating of the NMDA channel ([(*v* + 80)*/*60]^2^*/*[1 + ((*v* + 80)*/*60)^2^]).

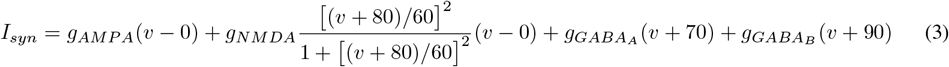

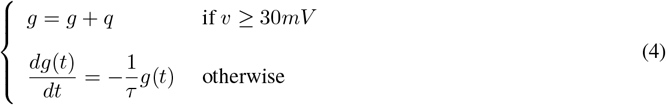

where the reversal potentials of AMPA and NMDA channels are 0 mV and reversal potentials for GABA_A_ and GABA_B_ channels are -70 mV and -90 mV respectively. The ratio of NMDA to AMPA receptors was set to 5:5, and the ratio of GABA_A_ to GABA_B_ receptors was set to 9:1 for all of the neurons in CA3 (105). For simplification, the synapse models of neurons in the DG included only AMPA and GABA_A_ receptors because learning does not occur in this region of the model (Table 2).

**Table 2:** Parameters for neurons organizing each layer.

| | N | $\tau_{AMPA}$ | $\tau_{NMDA}$ | $\tau_{GABA_A}$ | $\tau_{GABA_B}$ | $W_{Be}$ | $W_{Bi}$ |
| --- | --- | --- | --- | --- | --- | --- | --- |
| superficial EC | 16 | 5 | - | - | - | 5 | - |
| deep EC | 16 | 5 | - | - | - | 5 | - |
| Hilus(Exc) | 100 | 5 | - | 15 | - | 5 | 10 |
| Hilus(Inh) | 16 | 5 | - | - | - | 5 | - |
| GCL(Exc) | 800 | 5 | - | 15 | - | 5 | 10 |
| GCL(Inh) | 400 | 5 | - | - | - | 7 | 7 |
| CA3(Exc) | 2400 | 5 | 30 | 8 | 30 | 4, 0.75, 20 | 10 |
| CA3(Inh) | 120 | 5 | 30 | 8 | 30 | 5 | 10 |
| CA1 | 200 | 5 | 30 | - | - | 4 | - |

Synaptic conductance decreases exponentially with different time constants and increases by the peak conductance *q* upon the arrival of an action potential at the synapse (106). For connections without learning, the peak conductance *q* is initially set to 3 nS. For connections with learning, peak conductance *q* is initially set to 0 nS except for excitatory to inhibitory connections in CA3, where *q* is set to 0.5 nS to establish global inhibition as the default state, as detailed in the next section.

The total excitatory and inhibitory synaptic currents were limited to maximum values of *W*_*Be*_ and *W*_*Bi*_ respectively (Table 2). For excitatory neurons in CA3, the total synaptic current was differentiated based on the region of origin of the sending neuron—superficial EC, DG, or CA3—to preserve the distinct influences of each region. Additionally, some connections featured delayed transmission times to the postsynaptic neuron to ensure stable learning (Table 3).

**Table 3:** Parameters for connections between layers. Activation Phase indicates the connection status: E represents activation during the encoding phase, R during the retrieval phase, and A for activation across all phases. Physical dimension for *t*_*d*_ is ms.

| | $p_{conn}$ | $W$ | $t_d$ | Activation Phase | Learning |
| --- | --- | --- | --- | --- | --- |
| Superficial EC→Hilus (Exc) | 0.3125 | 3 | - | E | - |
| Superficial EC→Hilus (Inh) | One-to-One | 3 | - | E | - |
| Superficial EC→GCL (Exc) | 0.0625 | 2 | 10 | E | - |
| Superficial EC→GCL (Inh) | 0.5 | 4 | - | E | - |
| Superficial EC→CA3 (Exc) | 0.25 | 1 | - | R | O |
| Superficial EC→Deep EC | One-to-One | 3 | - | E | - |
| Deep EC→CA1 | 0.125 | 0.5 | - | E | - |
| Hilus (Inh)→Hilus (Exc) | 0.0625 | 4 | - | E | - |
| Hilus (Exc)→GCL (Inh) | 0.5 | 4 | - | E | - |
| GCL (Inh)→GCL (Exc) | 0.0013 | 3 | - | E | - |
| GCL (Exc)→CA3 (Exc) | 0.0125 | 2 | 3 | E | - |
| GCL (Exc)→CA3 (Inh) | 0.025 | 2 | - | E | - |
| CA3 (Exc)→CA3 (Exc) | 0.25 | 1 | 5 | R | O |
| CA3 (Exc)→CA3 (Inh) | 0.25 | 1 | - | R | O |
| CA3 (Exc)→CA1 | 0.5 | 0.15 | - | R | O |
| CA3 (Inh)→CA3 (Exc) | 0.25 | 2 | - | A | - |
| CA3 (Inh)→CA3 (Inh) | 0.167 | 0.5 | - | A | - |
| CA1→Deep EC | 0.125 | 2 | 15 | R | - |
| Noise→CA3 (Exc) | - | 0.5 | - | R | - |

### 5.4 STDP learning and heterosynaptic plasticity

Learning was implemented using symmetric STDP (55; 57). Instead of directly modifying the synaptic weights *W*, we adjusted the peak conductance *q* (106). Synaptic weights were initialized and kept constant throughout the learning process to maintain the network’s original structure and stability (Table 3). This approach ensures that modifications introduced by STDP refine existing connections without altering overall connection probabilities.

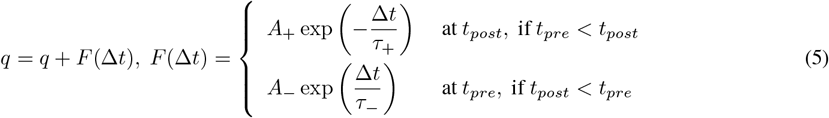

where Δ*t* = *t*_*post*_ − *t*_*pre*_ represents the time difference between action potentials, and *A*_±_ determines the maximum amount of synaptic modification, which decays exponentially with time constants *τ*_±_. Given the symmetric nature of the adopted STDP rule, *A*_±_ was set to 0.2 nS, and *τ*_±_ was set to 62.5 ms. Similar to general synaptic plasticity, when the time window between presynaptic and postsynaptic spikes is sufficiently short, the peak conductance is incremented by *F* (Δ*t*). However, with the adoption of heterosynaptic plasticity, the presynaptic stimulus no longer directly influences postsynaptic firing. The peak conductance is constrained within the range [0, *q*_*max*_], where *q*_*max*_ = 3 nS. The connections where learning occurs are listed in the Table 3.

### 5.5 Similarity analysis

Our model focuses exclusively on spatial input patterns and spatial pattern separation. Therefore, the similarity between firing patterns was assessed by calculating the spatial overlap, as suggested by (86).

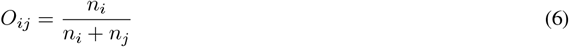

where *n*_*i*_ represents the number of activated neurons in pattern *i*, and *n*_*ij*_ denotes the number of neurons activated in both patterns *i* and *j*. The discrimination value Δ between patterns *i* and *j* is calculated as 1 − *O*_*ij*_. The firing pattern is determined based on neuron activation, except for the case of CA3. For CA3 patterns during retrieval, only neurons firing at rates exceeding 25 Hz are considered, ensuring that only significant activity is captured and noise is effectively filtered out. This threshold helps to isolate meaningful firing patterns that contribute to memory retrieval, thereby enhancing the accuracy of the pattern analysis.

## 6 Acknowledgements

The work was supported by the National Research Foundation of Korea (2022R1A2C1092831).

## 7 Author contributions

Gyeongtae Kim, Conceptualization, Investigation, Methodology, Software, Formal analysis, Visualization, Writing - original draft, Writing - review and editing; Pilwon Kim, Funding acquisition, Supervision, Writing - review and editing

## Notes

https://github.com/kgt1220/Hippocampus_SNN

### Competing Interest Statement

The authors have declared no competing interest.

### Summary of Updates

I revised the section explaining the model's architecture and operation to improve clarity.

